# KuafuPrimer: Machine learning empowers the design of 16S amplicon sequencing primers toward minimal bias for bacterial communities

**DOI:** 10.64898/2026.03.29.714677

**Authors:** Haoyu Zhang, Xiaoqing Jiang, Xiongwu Yu, Hongyi Wang, Ping Lu, Jiaheng Hou, Qian Guo, Tingting Xiao, Shufang Wu, Hengchuang Yin, Peter X. Geng, Jinyuan Guo, Alexandre Jousset, Zhong Wei, Yonghong Xiao, Huaiqiu Zhu

**Affiliations:** Department of Biomedical Engineering, and Department of Big Data and Biomedical AI, College of Future Technology, and Center for Quantitative Biology, Peking University, Beijing 100871, China; State Key Laboratory for Diagnosis and Treatment of Infectious Diseases, and The First Affiliated Hospital, Zhejiang University School of Medicine, Hangzhou 310003, China; Jiangsu Provincial Key Lab for Organic Solid Waste Utilization, Jiangsu Collaborative Innovation Center for Solid Organic Waste Resource Utilization, National Engineering Research Center for Organic-based Fertilizers, Nanjing Agricultural University, Nanjing 210095, China; Center of Artificial Intelligence for Digital Life, Institute for Artificial Intelligence, Peking University, Beijing 100871, China

**Author notes:** Corresponding author: Huaiqiu Zhu; Yonghong Xiao; Xiaoqing Jiang. These authors have contributed equally to this work and share first authorship (Haoyu Zhang, Xiaoqing Jiang).

## Abstract

Amplicon sequencing protocol targeting the 16S rRNA gene is a widely used and cost-effective method for exploring bacterial communities. However, its performance is often limited by primer bias arising from the arbitrary use of universal primers across diverse microbial communities and habitats. We propose KuafuPrimer to design the optimal 16S rRNA gene primers toward minimal bias for targeted bacterial communities, using few-shot machine learning to guide the primer design procedure based on a small number of samples. Simulations on 809 samples across 26 representative environments and habitats showed that KuafuPrimer-designed primers outperformed the universal primers in taxonomic accuracy, achieving an averaged 16.31% relative reduction in primer bias, with reductions up to 46.08% in plant samples. Notably, KuafuPrimer detected 29 rare and key taxa undetectable by the universal primers. Validation with 317 longitudinal gut microbiota samples demonstrated that KuafuPrimer-designed primers consistently outperformed the universal primers across temporal, individual, and cohort levels, with relative bias reductions of 5.03%, 3.53%, and 3.10%, respectively. Finally, in real PCR experiments on human gut samples from *Clostridioides difficile*-infected and healthy groups showed that polymerase chain reaction products using KuafuPrimer-designed primers correlated better with metagenomic data compared to the universal primers. More importantly, KuafuPrimer successfully detected *Clostridioides difficile*, the key pathogen missed by the universal primers, highlighting its potential for improving clinical diagnostics. In summary, KuafuPrimer provides a machine learning-based primer design strategy for targeted bacterial communities, with demonstrated utility in large-scale microbiome initiatives, longitudinal surveys and clinical diagnostics.

## Introduction

Microbes are ubiquitous and play essential roles in shaping human health, driving global biogeochemical cycles, and maintaining ecosystem balance^1–3^. The growing recognition of their significance has fueled an increasing demand for large-scale microbiome initiatives to construct comprehensive microbial maps and longitudinal surveys to monitor microbiota over time, which usually implies large-scale and laborious sequencing projects. Owing to its high cost-effectiveness, 16S rRNA gene amplicon sequencing has been preferred over metagenomics for microbiota exploration in large-scale microbiome initiatives (such as the Integrative Human Microbiome Project^4^ and the Isala Citizen-science Project^5^), longitudinal studies^6–10^, and consumer-oriented clinical diagnostics^11, 12^. However, most current 16S sequencing practices routinely apply universal primers without assessing their compatibility with the studied microbiota^13^, thereby introducing the well-known issue of “primer bias”. Primer bias is a technical limitation in which inappropriate primers used during targeted amplification lead to under- or over-representation of certain bacterial taxa, causing distorted estimates of community composition^14–17^ across various environmental and habitat-associated niches. For example, Anantharaman *et al.*^18^ discovered 46 novel phyla that were not detected by 16S sequencing with universal primers. The universal V1-V2 primer was reported to fail in detecting a few taxa, such as the health-promoting bacterium *Bifidobacteriales*^19, 20^, and the opportunistic pathogen *Fusobacteriota*^21^. Even the most commonly used V3 V4 primers have shown limited capability in identifying some genera, such as *Microbacterium*, *Tessaracoccus*, and *Opitutae*^22–24^, despite these genera being prevalent in various environments and habitats. Furthermore, some universal primers are prone to off-target amplification of host DNA, a major confounding factor in low microbial biomass samples, resulting in the loss of rare taxa, reduced bacterial resolution, and considerable sequence wastage^21, 25–27^. Primer bias largely affects the analyses of community structure, diversity, and bacterial population dynamics, resulting in the misinterpretation of microbiota, substantial waste of resources, and inaccuracies in diagnostics. It is attributed to several factors: (1) the polymorphisms in the conserved regions has led to a decline in the coverage rates of some primers^28^. (2) the taxonomic resolution of various V-regions varies in their capacity to amplify and classify specific lineages^29^, thereby precluding the design of a single optimal primer for diverse microbial habitats (3) the so-called universal primers for all microbiota, usually developed from a limited number of culturable taxa^16^, may offer insufficient coverage of many other community members.

This raises a key problem: how to develop 16S rRNA gene primers toward minimal bias to accurately characterize the studied microbial communities? Some efforts have been made to select primers tailored to specific environments^30, 31^. However, the key to addressing this problem lies in designing primers *ab initio* rather than selecting from an existing primer pool with a fixed list. Although several attempts to manually modify universal primers have been reported to largely improve their accuracy for certain environmental samples^21, 31–33^, the manual design of primers is labor-intensive and prone to errors^34^. Computational methods have started to be utilized for primer design in recent years^34–40^, but these methods encounter limitations when applied to designing optimal 16S rRNA gene primers for microbial communities. On one hand, these methods primarily develop primers based on a limited set of microorganisms, neglecting the heterogeneity and diversity of microbial communities, which results in poor performance in detecting underrepresented members. On the other hand, the 16S rRNA gene contains multiple V-regions interspersed with conserved regions, making current methods that rely on multiple sequence alignments (MSAs) of the full-length 16S rRNA gene both costly and imprecise^28, 41^. To sum up, there is an urgent need for an approach to design 16S rRNA gene primers *ab initio* that can account for the specific composition of target microbial communities and address the prevalent issue of primer bias.

Designing 16S rRNA gene primers toward minimal bias faces several challenges. The primary challenge lies in developing primers based on a small number of preliminary samples while ensuring their effectiveness on subsequent samples from relevant environments or habitats. This is essential for the integrative analysis of multicenter datasets, advancing personalized precision medicine, and exploring microbial dynamics. Machine learning offers a promising solution by training the model to capture the characteristics of microbial communities from a small set of samples. These characteristics have the potential to guide primer design, enabling the designed primers applicable to samples from related environments and habitats. The second challenge is addressing the limitations of existing design methods that rely on MSAs of the full-length 16S rRNA gene. Annotating the 16S rRNA gene represents a promising approach, enabling the MSA of conserved regions in primer design. However, current alignment-based approaches have proven ineffective for certain taxa, as reported in previous studies^29, 42^. Furthermore, alignment-free methods are also limited, as showed by our analysis, failing to annotate as much as 73,023 out of 238,890 (30.57%) 16S rRNA gene sequences within the SILVA dataset. Third, microbial communities are highly diverse and typically contain hundreds or even thousands of rare taxa. This “rare biosphere” represents a vast reservoir of genetic traits and specific ecological functions^43^, which deserves more focused attention.

To address these challenges, we develop KuafuPrimer, a machine learning-based method that recognizes community characteristics from preliminary samples to design 16S rRNA gene primers toward minimal bias for microbial communities (Fig. 1). To eliminate the reliance on MSAs of full-length 16S rRNA genes in the current primer design process, KuafuPrimer incorporates DeepAnno16, a deep learning–based algorithm for 16S rRNA gene annotation also developed in this study. We then adopt a few-shot machine learning strategy that trains the model on a small number of preliminary metagenomic samples from the target environments and habitats or hosts. Both simulations across different environments and real PCR experiments suggest that KuafuPrimer holds promise for advancing the rapid and cost-effective exploration of bacterial communities with minimal primer bias. This approach provides a general framework for the *ab initio* design of 16S rRNA gene primers tailored to target specific microbial communities.

**Figure 1.**
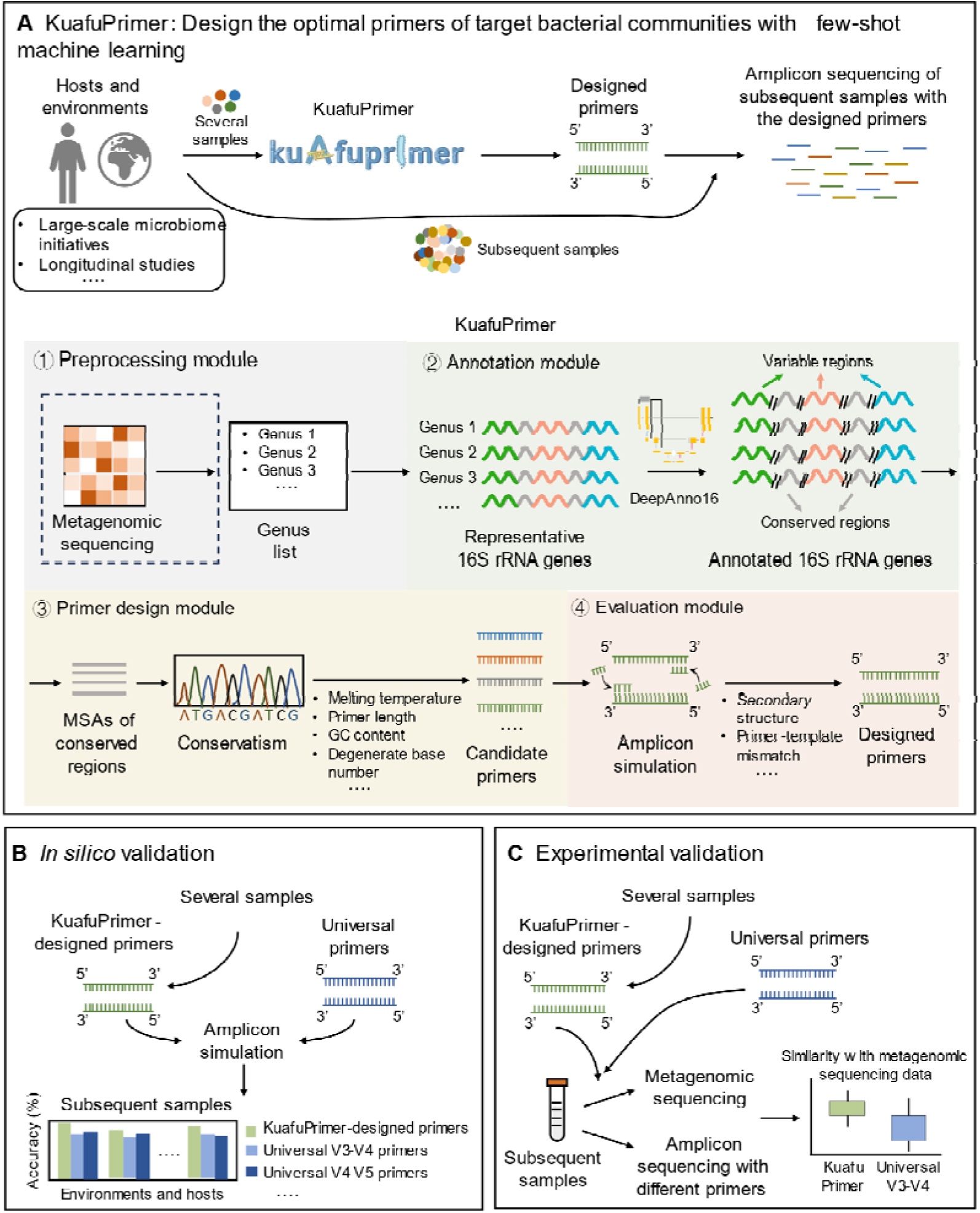
Schematic of workflow of KuafuPrimer. **A**. The workflow and usage paradigm of KuafuPrimer. KuafuPrimer designs minimal-bias 16S rRNA gene primers based on a small number of preliminary samples, making them applicable to subsequent samples from related environments and habitats. It consists of four modules: a preprocessing module, an annotation module, a primer design module, and an evaluation module. The *in silico* (**B**) and experimental (**C**) validation of KuafuPrimer.

## Results

### Machine learning empowered KuafuPrimer in primer design

For training KuafuPrimer to learn compositional characteristics (Fig. 2A), it takes as input a list of potential genera in the target microbial communities implying for a specific habitat, which can be generated either from a small number of metagenomic samples from relevant communities or simply based on prior knowledge. KuafuPrimer is composed of four modules: the preprocessing module, the annotation module, the primer design module, and the evaluation module. The preprocessing module provides representative sequences of the 16S rRNA gene from the input genus list. In the annotation module, KuafuPrimer introduces DeepAnno16 (Extended Data Fig. 1), a deep learning-based algorithm for rapid and efficient annotation of 16S sequences. The primer design module performs MSAs on the conserved regions to design all feasible primers targeting each possible V-region, while satisfying constraints about primer-template mismatch, primer length, GC content, secondary structure, and melting temperature^40, 44^. The evaluation module simulates the amplicons of each candidate primer pair for the target bacterial communities, assigns them to the genus level, and then evaluates the primer bias using the average taxonomic assignment accuracy. Finally, KuafuPrimer outputs primers toward minimal bias for the target microbial communities.

**Figure 2.**
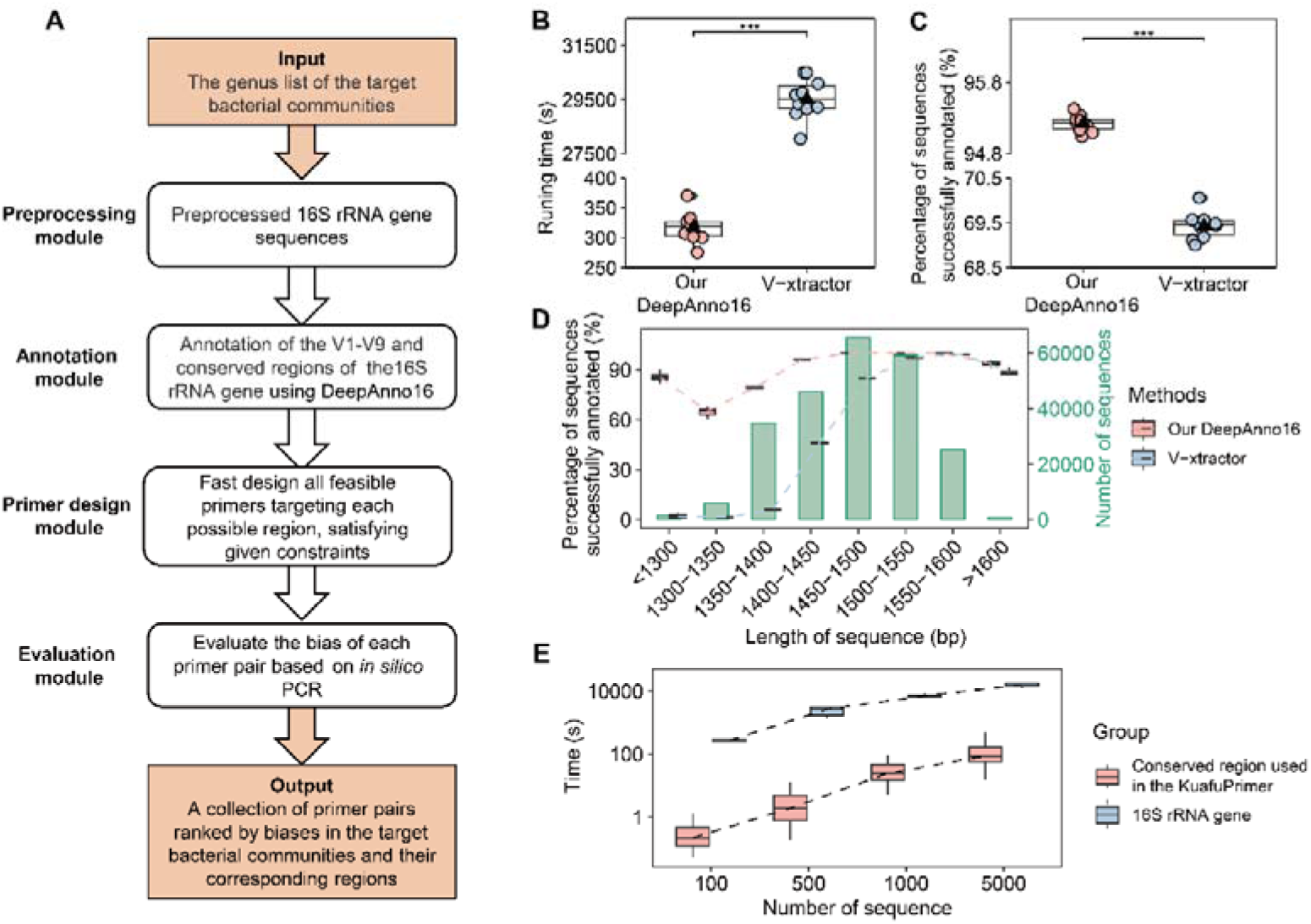
Overview of KuafuPrimer and performance of DeepAnno16. **A.** A more detailed workflow and description of the four modules of KuafuPrimer are provided. **B**. Comparison of running time between DeepAnno16 and V-xtractor. Comparison of the percentage of sequences successfully annotated by DeepAnno16 and V-xtractor (**C**) across varying sequence lengths (**D**). The Y-axis for the bar plot is positioned on the right. **E**. Running time comparison for MSAs using either the full-length 16S rRNA gene, or the conserved regions identified by DeepAnno16 and used in the primer design module of KuafuPrimer. ***: *p* < 0.001, **: *p* < 0.01, *: *p* < 0.05, Wilcoxon signed-rank test.

To improve the annotation success rate of 16S rRNA genes, we developed a deep-learning algorithm named DeepAnno16. In this study, we employed 10-fold cross-validation on a dataset comprising 238,890 16S rRNA gene sequences (see Methods) to train and assess its performance. We compared the performance of DeepAnno16 with V-xtractor^45^, the only available alignment-free tool for demarcating 16S rRNA gene sequences. As a result, DeepAnno16 successfully annotated all nine V-regions and conserved regions for 95.22 ± 0.11 % of 16S rRNA gene sequences, a large improvement over V-xtractor’s 69.43 ± 0.30 % (Fig. 2C, Wilcoxon signed-rank test, *p* < 0.001). In detail, compared to V-xtractor, DeepAnno16 had a higher success rate for the annotation of 16S rRNA gene sequences across various lengths, especially for those shorter than 1500 nucleotides, which account for 64.35 % of the dataset (Fig. 2D). This is mainly due to V-xtractor’s poor handling of the V1 and V9 regions located at the ends of 16S rRNA gene. Specifically, DeepAnno16 demonstrated significantly lower failure rates in annotating the V1 region, at 0.0080 ± 0.025 %, compared to 13.30 ± 0.26 % of V-xtractor. For the V9 region, DeepAnno16 exhibited failure rates of 4.76 ± 0.11 %, significantly lower than the 25.79 ± 0.21 % of V-xtractor (Extended Data Fig. 2A, Wilcoxon signed-rank test, *p* < 0.001). Moreover, DeepAnno16 exhibited a significantly shorter running time than V-xtractor (Fig. 2B, DeepAnno16 *vs.* V-xtractor: 317.95 ± 24.99 seconds *vs*. 29525.92 ± 45.78 seconds, Wilcoxon rank-sum test, *p* < 0.001). Furthermore, the results revealed that compared to V-xtractor, the conserved regions annotated by DeepAnno16 achieved a lower entropy value, a common index of measuring the variety of sites and a lower entropy value indicates more conservatism (Extended Data Fig. 2B-H)^46^.

MSA is the most critical step in primer design and, as an inherently NP-complete problem, it is also the speed-determining step^47^, therefore compromising the performance of existing methods that require MSAs of the full-length 16S rRNA gene. KuafuPrimer overcomes this limitation by designing primers only through MSAs of the conserved regions, rather than the full-length 16S rRNA gene. Moreover, using conserved regions for MSA significantly reduces computational time compared to using nearly full-length 16S rRNA genes. Specifically, for 100 sequences, the running time decreases by a factor of 715.98, and for 1,000 sequences, it decreases by a factor of 231.15 (Fig. 2E).

### *In silico* PCR amplification demonstrated KuafuPrimer’s superiority over universal primers

Following a common practice in published studies^31^, we evaluated the performance of KuafuPrimer using taxonomic assignment accuracy, defined as the proportion of 16S rRNA genes successfully amplified and accurately classified in the simulated PCR using the corresponding primers. We downloaded 809 metagenomic samples from various environments and habitats, including human gut (n = 269), human oral cavities (n = 37), human skin (n = 45), animals (n = 107), plants (n = 24), non-saline water (n = 194), saline water (n = 30), soil (n = 54), and wastewater treatment plants (WWTP, n = 49) (Supplementary Table S1-S2). We further divided these samples into 26 groups based on their literature annotations. For each group, five metagenomic samples were randomly chosen as the training set for KuafuPrimer, while the remaining samples served as the validation set, and this procedure was repeated ten times (see Methods). Five metagenomic samples were determined to be sufficient as a training set, covering 70.84 ± 15.21% of the genera present in the corresponding samples and achieving near-saturated taxonomic assignment accuracy for the designed primers. The stable and consistent binding sites further confirmed the reliability and robustness of the design process (Extended Data Fig. 3, Supplementary Table S3).

To evaluate the off-target effects of KuafuPrimer-designed primers, we downloaded 61,134 complete human mitochondrial genome sequences from the MITOMAP database^48^ (see Methods). The alignment results showed no non-specific amplification of human mitochondrial DNA by the optimal KuafuPrimer-designed primers for all 26 groups (Supplementary Table S4 and S5). However, consistent with previous studies^21^, the universal V4 and V4-V5 and primers exhibited off-target effects. Similarly, alignment against host chloroplast reference genomes for plant-derived groups (FS and Kal) showed that KuafuPrimer-designed primers also exhibited no off-target effects on host chloroplast DNA while several universal primers (universal V3-V4, V4, V4-V5, and V6-V8 primers) displayed off-target amplification^49^.

*In silico* PCR results showed that primers designed by KuafuPrimer significantly improved average taxonomic accuracy compared to commonly used universal primers targeting different V-regions (Fig. 3A and Supplementary Table S4 and S5). In detail, the accuracy of KuafuPrimer-designed primers was 88.15 ± 0.019%, while universal primers showed lower accuracy: V1-V3 primer-1 (76.39 ± 1.88%), V1-V3 primer-2 (62.16 ± 2.25%), V3-V4 primer (85.84 ± 1.75%), V4 (84.22 ± 2.40%), V4-V5 (85.59 ± 1.44%), V5-V7 (78.70 ± 2.86%), and V6-V8 (83.67 ± 1.16%) (Wilcoxon signed-rank test, all *p* < 0.001). Overall, KuafuPrimer-designed primers achieved a 16.31 % relative reduction in primer bias relative to the best-performing universal primer (V3-V4 primer) across all environments or habitats. KuafuPrimer-designed primers outperformed all others in 22 of 26 groups. In FS (olive cultivars FS17, plants), LAM (lake Alinen Mustajärvi, non-saline water), LL (lake Loclat, non-saline water), and PO (Pocket, human oral cavities) groups, the relative reduction in primer bias of KuafuPrimer-designed primers compared with the best-performing universal primer was 46.08%, 21.61%, 21.21%, and 20.41%, respectively (Wilcoxon signed-rank test, all *p* < 0.001). The accuracy of KuafuPrimer-designed primers slightly lower in the HC (healthy control group, human gut) group, partly due to the low coverage of the genera of HC samples used for primer design. Expanding the learning sample size from five to ten increased the genus coverage of the HC group from 50.77% to 61.87%, improving accuracy from 86.30% to 87.72%, which was comparable to universal primers (Supplementary Table S6).

**Figure 3.**
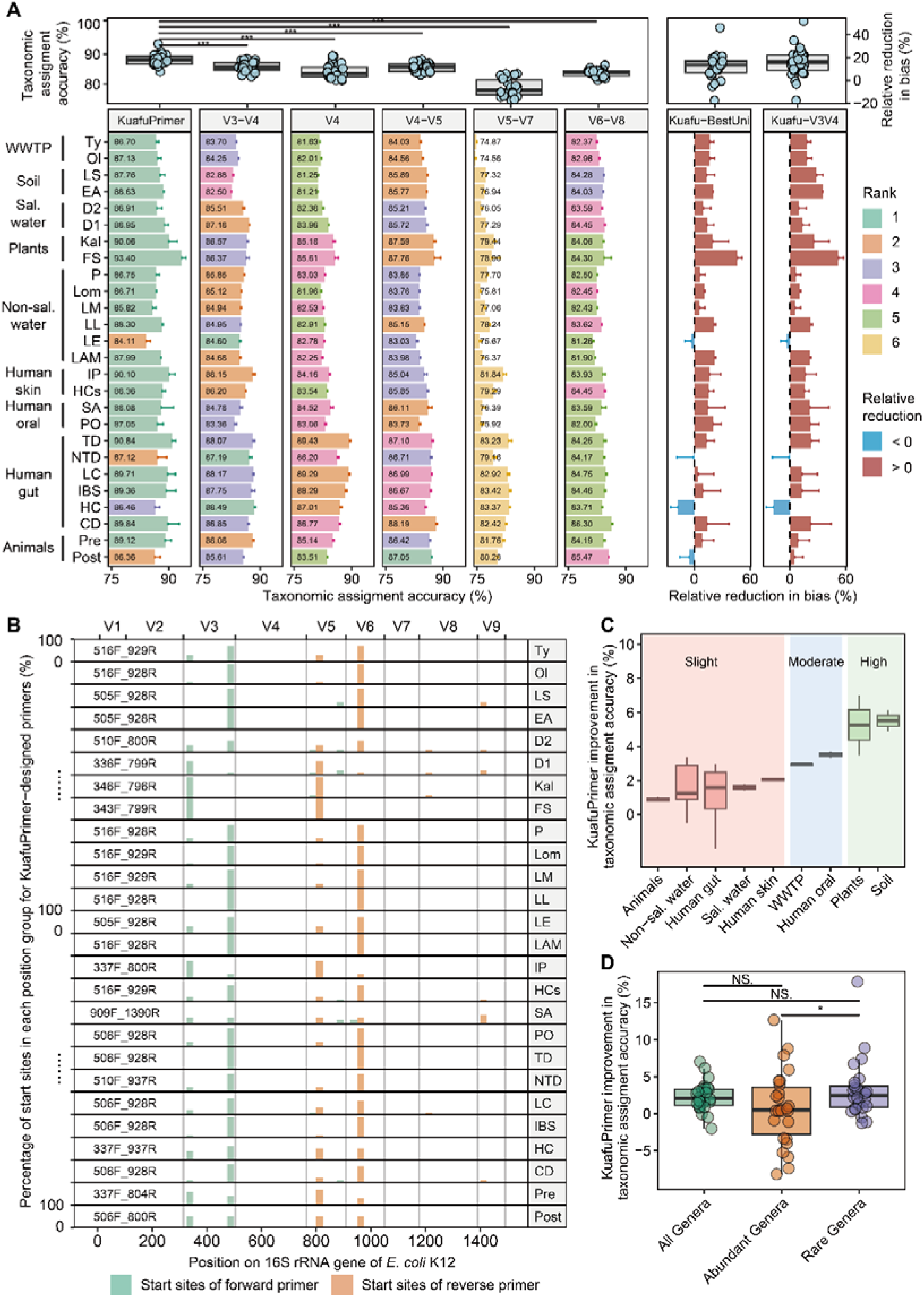
Comparison of *in silico* PCR simulations between KuafuPrimer-designed primers and various universal primers across 809 samples from a range of environments. **A.** Comparison of the taxonomic assignment accuracy of KuafuPrimer-designed primers and five universal primers. Detailed results including the other two universal primers are shown in Supplementary Table S5. Taxonomic assignment accuracy is defined as the proportion of 16S rRNA genes successfully amplified and accurately classified in the simulated PCR using the corresponding primers. The relative reduction in bias was defined as the decrease in bias (1 − taxonomic assignment accuracy) of KuafuPrimer-designed primers relative to the comparator primer, calculated as the percentage of the comparator’s bias, where the comparator was the best-performing universal primer for “Kuafu–BestUni” and the V3-V4 primers for “Kuafu–V3V4”. **B**. The distribution of target sites for the top-1 primers designed by KuafuPrimer in each group, with ten repetitions. **C**. Comparison of the taxonomic assignment accuracy improvement of KuafuPrimer over the V3-V4 primer in nine environments and habitats, categorized into slight, moderate, and high levels based on changepoint detection analysis. **D**. The average taxonomic assignment accuracy improvement of KuafuPrimer over the V3-V4 primer across all genera, abundant genera (relative abundance ≥ 1 %), and rare genera (relative abundance < 1 %) in each group. Sal water: saline water; Non-sal water: Non-saline water; Human oral: Human oral cavities. ***: *p* < 0.001, **: *p* < 0.01, *: *p* < 0.05, NS.: Not significant, Wilcoxon signed-rank test.

Through analyzing the target sites of the optimal primers designed by KuafuPrimer for each group, we discovered that these primers targeted different V-regions across samples from various environments and habitats, exhibiting high specificity for different microbial communities (Fig. 3B, Supplementary Table S5). In detail, for samples from soil, WWTP, and non-saline water, KuafuPrimer-designed primers mainly focused on the V4-V5 region, whereas for plant-derived samples, the focus shifted to the V3-V4 region. For samples from other environments or habitats, KuafuPrimer-designed primers always targeted at either the V3-V4 or V4-V5 regions, sometimes extending to the V6-V8 region. Notably, the optimal primer designed by KuafuPrimer for FS (olive cultivars FS17 group, plants), identified as 343F-799R, was positioned merely a few bp away from the universal V3-V4 primer (338F-806R) but showed a significant increase in taxonomic accuracy (Fig. 3A). These results indicate that KuafuPrimer is not only adept at identifying appropriate V-regions across a variety of environments and habitats but also in determining the optimal binding sites for primers targeting these V-regions. Moreover, the differences in targeted V-regions and binding sites of the optimal primers designed by Kuafuprimer for different environments further highlight the inherent challenge and infeasibility of creating a universally optimal primer with the best performance across all microbial communities.

We subsequently evaluated the taxonomic accuracy improvement of primers designed by KuafuPrimer compared to the universal V3-V4 primer. The latter was ranked second in simulation-based accuracy assessments (Fig. 3A) and was the preferred primer in numerous studies^50^. Changepoint detection analysis revealed that KuafuPrimer-designed primers significantly improved accuracy predominantly for soil and plant samples (Fig. 3C, 5.50 ± 0.88 % and 5.26 ± 2.51 %, respectively), followed by human oral cavities and WWTP samples (3.49 ± 0.28 % and 2.94 ± 0.093 %, respectively). Improvements were also observed in human skin, non-saline water, saline water, human gut, and animal samples, with respective increases of 2.06 ± 0.15 %, 1.59 ± 1.51 %, 1.58 ± 0.26 %, 1.14 ± 1.89 %, and 0.90 ± 0.21 %. These findings align with previous studies indicating that the universal V3-V4 primer is more suitable for samples from humans, animals, and non-saline water^20, 25, 30, 51, 52^, while its accuracy diminishes for samples from soil and plant environments^26, 50, 53^. These results highlight the necessity and effectiveness of KuafuPrimer in enabling the accurate detection of bacterial community structure and diversity.

Interestingly, we discovered that KuafuPrimer significantly improved taxonomic accuracy for rare genera compared to abundant genera (Fig. 3D). This may be because universal primers are predominantly developed from a limited number of cultured taxa^16^, whereas KuafuPrimer, by utilizing several metagenomic samples as training set, is capable of covering most microbial members present in the communities. Furthermore, *in silico*results showed that KuafuPrimer-designed primers outperformed the universal V3-V4 primer by at least 90 % in accuracy across 29 genera, despite lower performance in five genera (Fig. 4A and 4B). Among these 29 genera, KuafuPrimer-designed primers exhibited higher accuracy for eight genera in at least five different environments or habitats. Five of the eight genera, namely, *Microbacterium*^22, 54^, *Cutibacterium*^54, 55^, *Tessaracoccus*^23^, *Propionibacterium*^54, 55^, and *Agrococcus*^56^ have been consistently reported for their poor capture by the V3-V4 primer. These five genera, isolated from a wide variety of habitats, play essential roles in many processes, such as soil nutrient enhancement^57, 58^, plant growth promotion^59, 60^, skin health maintenance^61^, and heavy metal detoxification^58^, thus should not be overlooked. Additionally, some other key taxa were accurately identified by KuafuPrimer-designed primers, whereas they failed to be detected by the V3-V4 primer. These taxa included the short-chain fatty acid producers such as *Acidipropionibacterium*^62^, *Leuconostoc*^63^, and *Akkermansia*^64–66^ from human samples; the polyphosphate-accumulating bacterium *Microlunatus*^67^ and the organohalide-respiring bacterium *Dehalococcoides*^68^ from wastewater treatment plant samples; potential freshwater quality indicators like *Luteolibacter*^69^ and the aquatic probiotics *Roseimicrobium*^70^ from water samples; as well as *Nocardiopsis*, noted for its prolific production of natural products and environmental adaptability^71^. These findings suggest that, compared to the universal V3-V4 primer, KuafuPrimer-designed primers can also accurately identify many more key taxa that play critical roles in environmental and habitat samples.

**Figure 4.**
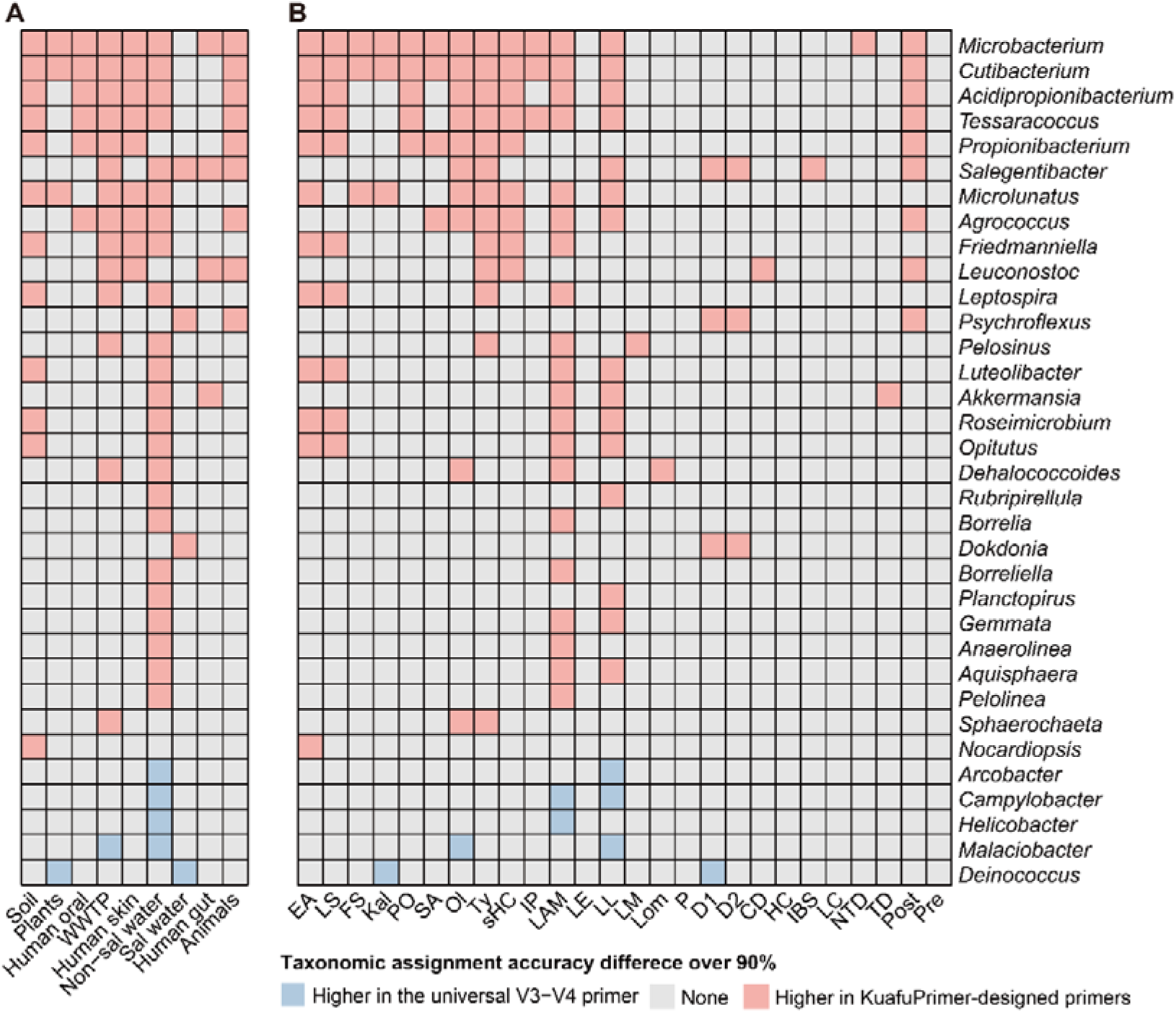
Genera with over a 90 % taxonomic assignment accuracy difference between KuafuPrimer-designed primers and the universal V3-V4 Primer. For 34 genera, the difference in taxonomic assignment accuracy between KuafuPrimer-designed primers and the universal V3-V4 primer exceeded 90 %. The heatmaps display which primer achieved higher accuracy for these 34 genera across nine environments (**A**) and 26 groups (**B**). Red: genera where primers designed by KuafuPrimer outperformed the universal V3-V4 primer in accuracy by ≥ 90 %. Blue: genera where the universal V3-V4 primer outperformed KuafuPrimer-designed primers in accuracy by ≥ 90 %. Grey: genera not detected in metagenomic samples or with an accuracy difference of less than 90 %.

### *In silico* PCR amplification of longitudinal samples demonstrated KuafuPrimer’s advantage across different time points, individuals, and cohorts

To assess whether primers designed by KuafuPrimer based on several early samples, herein usually meaning a small number of sample size for our few-shot learning strategy, could be applied to subsequent samples from the same individual, different individuals, and different cohorts, we downloaded 317 gut samples from 28 healthy individuals from three public studies (Supplementary Table S7). The sampling period for each individual lasted 9.71 ± 2.21 months, ranging from 7 to 12 months, with nearly one sample collected per month. Samples collected during the first two months from each individual served as the training set for primer design with KuafuPrimer. Subsequent samples, including those from the same individual, the same cohort, and different cohorts, were used for testing.

*In silico* PCR results demonstrated that KuafuPrimer-designed primers significantly increased average taxonomic assignment accuracy compared to several universal primers targeting various V-regions (Fig. 5A). Specifically, KuafuPrimer-designed primers achieved average taxonomic assignment accuracies of 77.93 ± 1.73%, 77.58 ± 1.22%, and 77.48 ± 1.18% for samples from across temporal, individual, and cohort levels, respectively. They were significantly higher than those obtained with universal primers: V1-V3 primer-1 (67.80 ± 0.82%), V1-V3 primer-2 (58.00 ± 0.79%), V3-V4 primer (76.76 ± 1.54%), V4 primer (72.02 ± 1.36%), V4-V5 primer (72.56 ± 1.12%), V5-V7 primer (65.08 ± 1.11%), and V6-V8 primer (73.53 ± 1.16%) (Wilcoxon signed-rank test, all *p* < 0.001). Compared to the best-performing universal V3-V4 primer, KuafuPrimer-designed primers achieved relative reductions in primer bias of 5.03%, 3.53%, and 3.10% for intra-individual, cross-individual, and cross-cohort samples, respectively. Additionally, no significant difference was observed in the average taxonomic assignment accuracy of KuafuPrimer-designed primers when applied to the same individuals, different individuals, or different cohorts, underscoring the comparable applicability of primers designed by KuafuPrimer based on several early samples for subsequent relevant samples.

**Figure 5.**
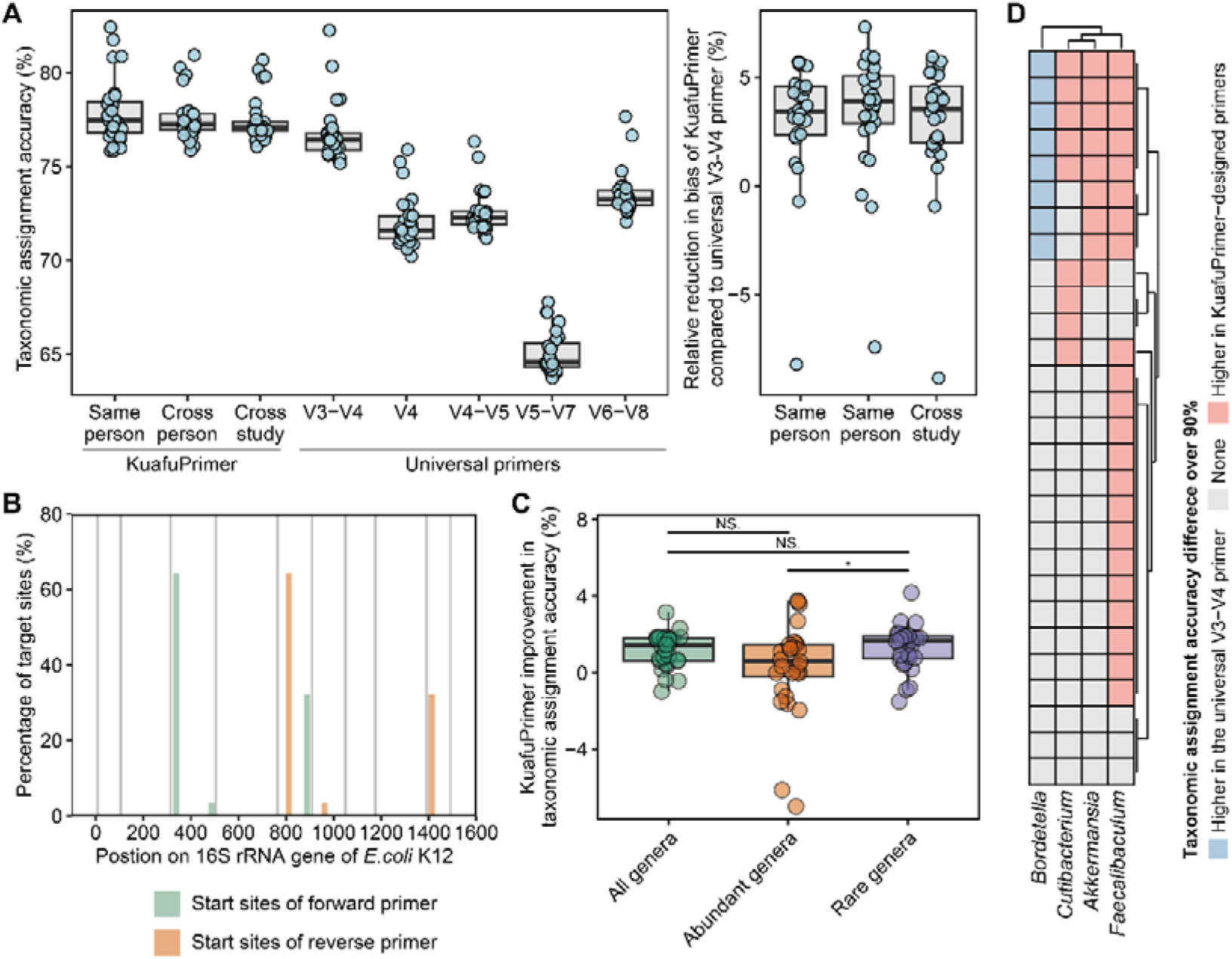
Performance of *in silico* PCR simulations of KuafuPrimer-designed primers in longitudinal studies. **A**. Comparison of the taxonomic assignment accuracy of KuafuPrimer-designed primers with five universal primers. Detailed results for the other two universal primers are provided in Supplementary Table S8. Taxonomic assignment accuracy is defined as the proportion of 16S rRNA genes successfully amplified and accurately classified in simulated PCR using the corresponding primers. All pairwise group comparisons are significantly different, except for those marked “NS” (not significant). The relative reduction in bias was calculated as the decrease in bias (1 − taxonomic assignment accuracy) achieved by KuafuPrimer-designed primers compared to the V3-V4 primer, expressed as a percentage of the V3-V4 primer’s bias. **B**. Distribution of target sites for the top-1 primers designed by KuafuPrimer for each subject. **C**. Average improvement in taxonomic assignment accuracy of KuafuPrimer-designed primers over the V3-V4 primer across all genera, abundant genera (relative abundance ≥ 1%), and rare genera (relative abundance < 1%) for each subject. *: p < 0.05, NS: Not significant (Wilcoxon signed-rank test). D. For four genera, the difference in taxonomic assignment accuracy between KuafuPrimer-designed primers and the universal V3-V4 primer exceeded 90 %. The heatmaps show which primer achieved higher accuracy for these four genera across individual subjects. Red: genera where KuafuPrimer-designed primers outperformed the universal V3-V4 primer by ≥ 90 % taxonomic assignment accuracy. Blue: genera where the universal V3-V4 primer outperformed KuafuPrimer-designed primers by ≥ 90 % taxonomic assignment accuracy. Grey: genera either not detected in metagenomic samples or with a taxonomic assignment accuracy difference of less than 90 %.

Further investigation of the target regions of KuafuPrimer-designed primers revealed that, while the optimal primers for 62.39% (18/28) of individuals targeted the V3-V4 region, 32.14% (9/28) targeted the V6-V8 region, and 3.58% (1/28) targeted the V4-V5 region (Fig. 5B). This suggests that KuafuPrimer is able to design optimal primers for different individuals through a comprehensive consideration of all candidate V-regions. Next, we compared KuafuPrimer with the universal V3-V4 primer, which is frequently used in studies of the human gut microbiome, such as the Human Microbiome Project^72^, and which ranked the highest among all universal primers in our *in silico* PCR results. Alignment with 61,134 human DNA sequences revealed that both the KuafuPrimer-designed primers and the V3-V4 primer exhibited no off-target effects in these cases (Supplementary Table S8). Once again, we found that KuafuPrimer significantly enhanced taxonomic assignment accuracy for rare genera, rather than abundant genera (Fig. 5C, Wilcoxon rank-sum test, *p* < 0.001), when compared to the universal V3-V4 primers. Moreover, *in silico* PCR results showed that KuafuPrimer-designed primers outperformed the universal V3-V4 primer by at least 90% in accuracy across three genera, including short-chain fatty acid producers *Akkermansia*^64–66^ and *Faecalibaculum*^73^, despite lower performance in one genus (Fig. 5D). Thus, by delivering higher performance in taxonomic assignment accuracy and the identification of key and rare microbial members within individual microbiomes, KuafuPrimer stands out in uncovering microbial dynamics and holds great promise for advancing personalized precision medicine.

### PCR experiments validated the superior performance of KuafuPrimer

*Clostridioides difficile* infection (CDI) is a leading cause of healthcare-associated infections, largely contributeing to morbidity and mortality in hospitalized patients^74^. To evaluate the applicability of KuafuPrimer in clinical diagnostics, we collected 19 fecal samples during 2018–2019, categorizing them into CDI positive (CD+, n = 12) and CDI negative (CD-, n = 7) groups (see Methods and Supplementary Table S10). Using KuafuPrimer, we automatically designed minimal-bias primers based on these preliminary samples and applied them to 34 fecal samples collected in 2023, consisting of 17 CD+ and 17 CD− samples (Fig. 6A). Furthermore, we characterized microbial community composition through both 16S rRNA gene sequencing using the universal V3-V4 primer and shotgun metagenomic sequencing. Alignment of PCR results generated by KuafuPrimer-designed primers and universal primers with the human genome revealed no matches, indicating the absence of off-target effects. We then compared the PCR products obtained with KuafuPrimer-designed primers against the results from the universal V3-V4 primer and metagenomic sequencing.

**Figure 6.**
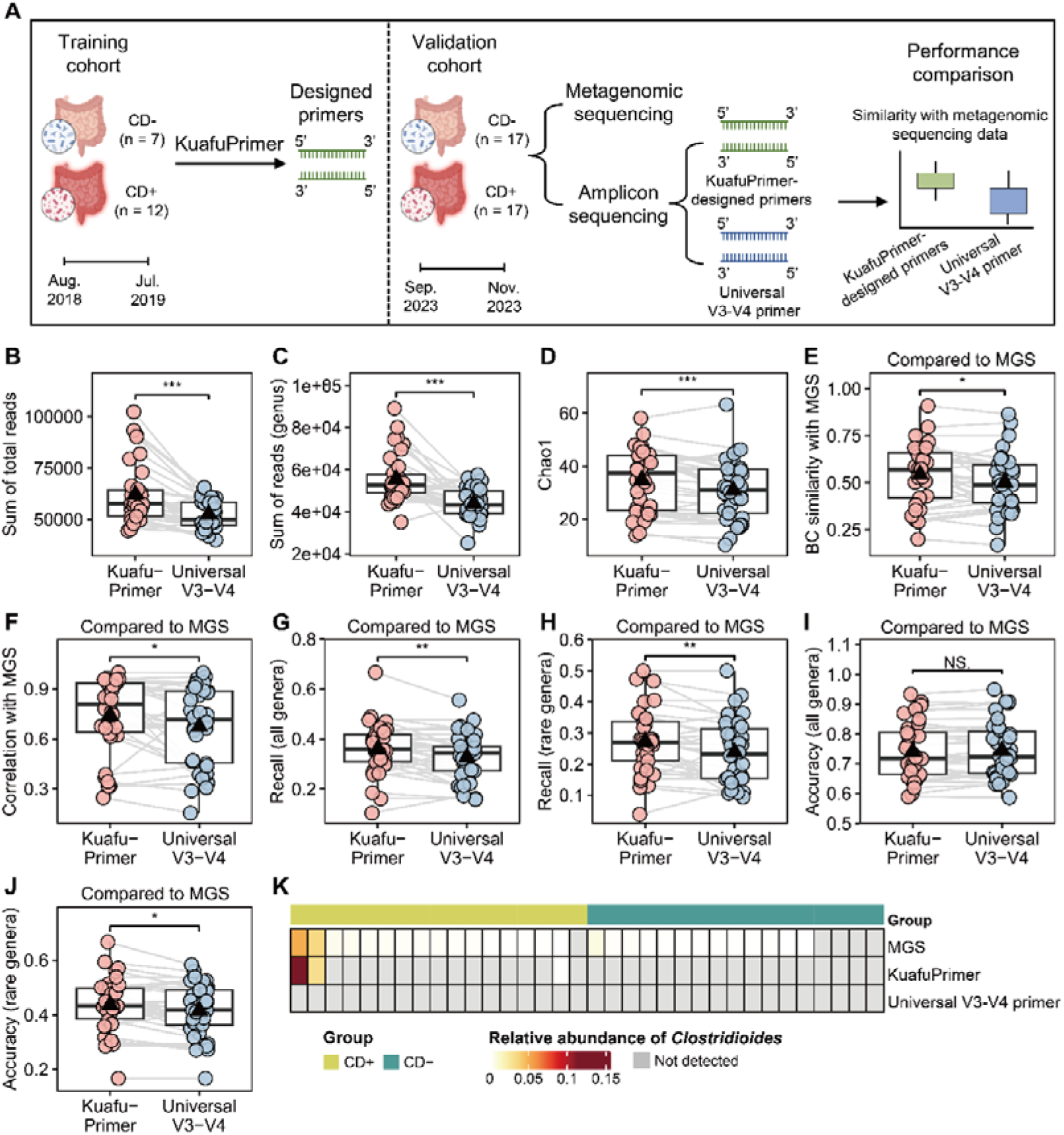
Comparison of PCR experimental results between KuafuPrimer and the universal V3-V4 primer in human gut samples. **A.** The workflow of experimental validation of KuafuPrimer. Comparison between PCR products of KuafuPrimer-designed primers and the universal V3-V4 primer in human gut samples in terms of the total reads (**B**), reads assignment to the genus level (**C)**, and Chao1 index (**D**). We further evaluated the performance of KuafuPrimer-designed primers and the universal V3–V4 primer by comparing them with metagenomic sequencing (MGS) data. Specifically, we assessed Bray–Curtis similarities (**E**) and Pearson correlation coefficients of their PCR products with results of MGS (**F**). Using the genera identified by MGS as the gold standard, we further calculated the recall of all genera (**G**), recall of rare genera (**H**), accuracy of all genera (**I**), and accuracy of rare genera (**J**) for both KuafuPrimer-designed primers and the universal V3-V4 primer. **K**. Relative abundance of *Clostridioides* detected by MGS, KuafuPrimer-designed primer, and the universal V3-V4 primer. BC: Bray-Curtis. MGS: Shotgun metagenomic sequencing. ***: *p* < 0.001, **: *p* < 0.01, *: *p* < 0.05, NS.: Not significant, Wilcoxon signed-rank test. CD+: CDI positive groups, n = 17; CD-: CDI negative group, n = 17. Part of the figure was created with BioRender.com.

The experimental results showed that, compared to the V3-V4 primer, KuafuPrimer-designed primers yielded a significantly higher total reads count (Fig. 6B, KuafuPrimer: 62211.18 ± 14645.87, V3-V4: 52554.47 ± 7067.49; Wilcoxon signed-rank test, *p* < 0.001) and a greater number of reads classified at the genus level (Fig. 6C, KuafuPrimer: 55543.18 ± 11429.94, V3-V4: 44026 ± 6832.44; Wilcoxon signed-rank test, *p* < 0.001). To alleviate the potential influence of sequencing depth variation, we rarefied the genus table to the smallest sequence count observed across all samples. The relative abundances of genera obtained using KuafuPrimer-designed primers and the universal V3-V4 primer are shown in Extended Data Fig. 4A. We conducted a differential analysis comparing the relative abundance of genera acquired by KuafuPrimer and the universal V3-V4 primer. We discovered that KuafuPrimer effectively identified some genera that weren’t detected by the universal V3-V4 primer, including *Microbacterium*, *Cutibacterium*, *Tessaracoccus*, *Propionibacterium*, *Agrococcus*, *Borrelia*, and *Dokdonia*. These genera were also consistently reported in multiple studies as either underrepresented or undetected when using the universal V3-V4 primer^22–24, 54, 56^. Additionally, compared to the universal V3-V4 primer, KuafuPrimer-designed primers retrieved a higher genus richness, as indicated by the Chao1 index (Fig. 4D, KuafuPrimer: 34.89 ± 11.50, V3-V4: 30.94 ± 11.13; Wilcoxon signed-rank test, *p* < 0.001). These experimental results demonstrate that KuafuPrimer-designed primers outperform the universal primers in terms of obtaining more reads, retrieving higher richness, and detecting rare taxa.

PCR results utilizing primers designed by KuafuPrimer were more similar to metagenomic data than those obtained with the universal V3-V4 primer, demonstrated by significantly higher Bray-Curtis similarity (Fig. 6E, KuafuPrimer: 0.54 ± 0.16, V3-V4: 0.50 ± 0.16; Wilcoxon signed-rank test, *p* < 0.05) and Pearson’s correlation coefficients (Fig. 6F, KuafuPrimer: 0.74 ± 0.22, V3-V4: 0.67 ± 0.24; Wilcoxon signed-rank test, *p* < 0.05) with metagenomic samples. Furthermore, primers designed by KuafuPrimer also exhibited a higher recall, as evidenced by a higher proportion of genera present in metagenomic samples that could be successfully amplified using KuafuPrimer-designed primers (Fig. 6G, KuafuPrimer: 0.36 ± 0.10, V3-V4: 0.33 ± 0.093; Wilcoxon signed-rank test, *p* < 0.01). Interestingly, compared to the V3-V4 primer, KuafuPrimer-designed primers showed a higher recall of rare taxa (Fig. 6H, KuafuPrimer: 0.27 ± 0.10, V3-V4: 0.24 ± 0.103; Wilcoxon signed-rank test, *p* < 0.01), rather than abundant genera (Extended Data Fig. 4B, Wilcoxon signed-rank test, *p* > 0.05). Moreover, KuafuPrimer-designed primers also showed higher accuracy in identifying rare taxa (Fig. 6J, KuafuPrimer: 0.44 ± 0.097, V3-V4: 0.42 ± 0.094; Wilcoxon signed-rank test, *p* < 0.05), while no significant difference was found in all genera and abundant genera (Fig. 6I, Extended Data Fig. 4C, Wilcoxon signed-rank test, *p* > 0.05). These results demonstrate that KuafuPrimer can design primers based on earlier samples and apply them to later ones, even those collected four years later and from different individuals, achieving smaller biases compared to universal primers.

It is noteworthy to mention that although *C. difficile* was detected through clinical CDI diagnosis in all samples from the CD+ group, the V3-V4 primer failed to capture any sequences belonging to *Clostridioides*. In contrast, KuafuPrimer-designed primers accurately identified *Clostridioides* in three samples within the CD+ group and detected no *Clostridioides* sequences in the CD− samples, highlighting KuafuPrimer’s high specificity and sensitivity in identifying *Clostridioides* (Fig. 6M). KuafuPrimer successfully detected *Clostridioides* in two metagenomic samples where its relative abundance exceeded 3%. However, KuafuPrimer failed in detecting *Clostridioides* in some other samples where its relative abundance was below 1%, likely due to the limitations of 16S rRNA gene sequencing methods in detecting genera with low relative abundance. This supports that KuafuPrimer-designed primers are effective in accurately identifying key taxa within studied communities.

## Discussion

The 16S amplicon sequencing remains the most commonly used technique for analyzing bacterial communities, as it is more cost-effective and convenient than whole-metagenome sequencing^75, 76^. However, primer bias in 16S rRNA gene sequencing is a recognized source of distortion that can lead to certain bacterial genera being underrepresented or entirely absent from estimated taxonomic profiles. Moreover, the varying capacity of different 16S rRNA gene variable regions to amplify and classify distinct taxonomic lineages, coupled with the accumulation of polymorphisms in conserved regions and the immense diversity of microbiota, makes it challenging to design a primer that is optimal for all bacterial communities. Thus, we introduced KuafuPrimer to enable *ab initio* design of 16S rRNA gene primers tailored for the studied bacterial communities. KuafuPrimer: (1) can reduce the primer bias *in silico* and experimentally, yielding more accurate community profiles and recovering rare or clinically critical taxa; (2) learns microbial characteristics from a small pilot set and designs primers that remain effective for subsequent samples from the relevant environment or habitat, (3) accelerates and improves primer design process by aligning only the short conserved binding sites, enabling exhaustive evaluation of every candidate primer across all hypervariable regions in one pass. Collectively, these features make KuafuPrime a rapid, low-bias alternative to universal primersthat can improve cohort-tailored 16S rRNA profiling while remaining compatible with standard amplicon workflows.

KuafuPrimer’s findings underscore that high-fidelity 16S-rRNA profiling is influenced by both the choice of hypervariable region and the exact primer-binding site. KuafuPrimer typically favored primers targeting the V3-V4 or V4-V5 regions, but for specific cohorts — saliva (Saliva group, human oral cavities) and loessal soil (Loessal Soil group, soil) — it suggested V6-V8 region as the optimal. Previous research has also highlighted the importance of frequently neglected V9 region, due to its efficacy in classifying genera such as *Staphylococcus* and *Clostridium*^29, 42^. *In silico* analyses of longitudinal gut microbiota revealed that primers targeting regions such as V6-V8 or V4-V5 can outperform the universal V3-V4 primers, and *in vivo* gut data showed that KuafuPrimer-designed primers targeting at V4-V5 region generated amplicon profiles more congruent with metagenomic sequencing. Even within the same hypervariable region, fine-scale refinement matters: in FS group (olive cultivars FS17 group, plants), switching the universal V3-V4 set (338F-806R) with KuafuPrimer’s optimized pair (343F-799R) increased taxonomic accuracy from 86.37 % to 93.40 %. Complementary research has likewise shown that altering a single nucleotide can markedly improve coverage of key genera such as *Bifidobacterium*^30^ and *Fusobacteriota*^21^.

Despite constituting a minor fraction of the total microbiota abundance, rare-biosphere bacteria offer a vast reservoir of ecological function and resilience, exhibiting specific and even unique ecological and biogeographical characteristics that can differ significantly from abundant bacteria^16^. Simulations based on 809 microbial samples suggest that KuafuPrimer achieves greater accuracy in identifying rare and key taxa, which are crucial for understanding environmental specificity. Experimental evidence further supports that, compared to the universal primer, KuafuPrimer-designed primers exhibit significantly better recall and accuracy for rare genera. Notably, KuafuPrimer successfully detect *C. difficile* in CD+ groups, which serves as the key taxon for the study, whereas the universal primer fails to identify it. The accuracy of KuafuPrimer in detecting rare and key taxa can be partly explained by its effective use of five metagenomic samples for training, achieving about 70 % coverage of the bacterial communities in the studied environments or habitats. Conversely, universal primers are predominantly designed from a limited number of cultured bacteria^16^. The rapid expansion of bacterial database entries further poses a considerable challenge for detecting rare taxa using universal primers, since most bacteria in these databases lack cultured representatives^77, 78^. In contrast, KuafuPrimer can be easily updated to incorporate these latest entries, thereby enhancing the amplification efficiency of the designed primers.

There are also some limitations in the KuafuPrimer developed in this study. For instance, in the evaluation module, it only considers constraints including primer length, GC content, secondary structure, mismatch count, and melting temperature. Other constrains such as Gibbs free energy of thermodynamics also play pivotal roles in the efficient annealing of primers and target^44^. Additionally, the impact of sequencing errors is not taken into account in this study. Future versions will incorporate additional thermodynamic parameters and evaluate robustness under realistic sequencing-error profiles using simulated data. Furthermore, KuafuPrimer is trained using full-length 16S rRNA gene sequences from the SILVA database. Extending KuafuPrimer to accommodate full-length 16S rRNA gene sequences de novo assembled from metagenomic data could broaden applicability and facilitate detection of novel taxa. Additionally, while our results provide a proof-of-concept for the value of KuafuPrimer, broader experimental validation across additional community types and study designs will further strengthen its generalizability.

## Methods

### Construction of Datasets

For DeepAnno16 training, we collected 431,388 non-redundant bacterial 16S rRNA gene sequences from SILVA SSU Ref NR 99 database (release 138.1)^79^– the most widely used high-quality ribosomal RNA reference in 16S studies. Nine hypervariable regions (V1-V9) and the conserved regions of the 16S rRNA gene were annotated as described in previous studies with primer contigs (Supplementary Table S9)^80^. Sequences with at least one primer mismatched or aligned in the wrong way were removed. As a result, 238,890 accurately annotated 16S rRNA gene sequences formed the training dataset for DeepAnno16.

To ensure that all representative 16S sequences included the integrated primer-binding regions across all hypervariable domains, the 238,890 16S rRNA gene sequences were then aligned against the reference sequence of *Escherichia coli* K-12 MG1655 (NCBI Gene ID: 947777). Among them, 145,137 full-length 16S rRNA gene sequences that covered the 8th to 1,492nd sites in *E. coli* were selected as the complete-core dataset for further primer design using KuafuPrimer and *in silico* evaluation of primer performance.

### KuafuPrimer algorithm

The workflow of KuafuPrimer consists of four major modules (Fig. 2A), illustrated as follows:

#### Preprocessing module

The relevant genus list based on the analysis of several metagenomic samples from relevant communities or the prior knowledge of the target environmental communities is the only input required for KuafuPrimer. A certain amount (default: 100) of 16S rRNA gene sequences for each genus are collected from the constructed dataset as the representative sequences in the target communities for primer design.

#### Annotation module

To annotate all candidate hypervariable regions and extract associated primer-binding regions from the relevant representative sequences, we construct a deep learning module named DeepAnno16. DeepAnno16 is built upon a SE-ResNet based encoder-decoder deep neural network comprising interconnected down-sampling and up-sampling blocks, inspired by the widely used U-net architecture in the semantic segmentation field (Extended Data Fig. 1)^81^. DeepAnno16 was trained using a 10-fold cross-validation strategy, with dropout and early stop strategies employed to reduce overfitting (detailed description of DeepAnno16 in Supplementary methods).

#### Primer design module

To design forward and reverse primers for each candidate combination of V-regions, Muscle (version 5.1)^82^ is used to conduct MSAs for the associated set of fragments of conserved sub-region extracted by the annotation module. Afterward, all feasible primers satisfying constraints of primer length, GC content, and melting temperature with at most three degenerate bases are designed based on MSAs using homemade Python scripts (Supplementary methods).

#### Evaluation module

All possible combinations of forward and reverse primers with a melting temperature difference of 5 C or lower constitute the candidate sets of primer pairs. MFEprimer (version 3.2.6)^83^ is employed to remove primer pairs that may produce dimer or hairpin secondary structures. We employ an *in silico* PCR process to evaluate the primer bias of each primer pair for the studied bacterial communities. First, BLAST (version 2.6.0+) is used to obtain the simulated amplicons of all representative sequences. The amplification process is considered successful if there is no mismatch between primer and template. Subsequently, some commonly used 16S rRNA gene classifiers are used to taxonomically assign the simulated amplicons according to the SILVA v138.1 nr database (default: RDP classifier in Mothur version 1.48.0). Finally, a collection of primer pairs ranked by taxonomic accuracy at the genus level (the proportion of representative sequences accurately assigned for each genus) and their corresponding target hypervariable regions are outputted. The evaluation of off-target amplification of primers is conducted using the BLAST tool to align them against the corresponding off-target sequence database (default: the MITOMAP database^48^). A primer is considered to amplify an off-target sequence when there is no mismatch between them.

KuafuPrimer is developed in Python 3.8. It is noteworthy that parameters and some external tools used herein can be customized by users based on their usage circumstances, and the default configurations are utilized in this study.

### Collection of public metagenomic samples

Metagenomic samples were collected from several published studies (Supplementary Tables S1, S2, and S7). Only paired-end sequencing samples generated using the Illumina platform with at least 1,000,000 raw reads were included. Only groups with more than 10 samples were retained, yielding a total of 809 samples across 26 groups, including human gut (n = 269), human oral cavities (n = 37), human skin (n = 45), animals (n = 107), plants (n = 24), non-saline water (n = 194), saline water (n = 30), soil (n = 54), and wastewater treatment plants (WWTP, n = 49). From each group, 5 samples were randomly selected for training, while the remaining samples were used for testing, with the process repeated 10 times. For longitudinal studies, only subjects with more than 6 samples and a sampling duration exceeding 6 months were included, resulting in 317 samples across 28 subjects. For each subject, samples from the first two months (at least 2 samples) were used for training, while samples from subsequent time points (from the same or different subjects) were used for testing.

### Recruitment of Subjects for CDI samples

#### Training cohort

A cohort of 19 stool samples was collected from patients newly admitted to the intensive care units between August 2018 and July 2019 at the First Affiliated Hospital of Zhejiang University School of Medicine in Hangzhou, China. These samples were divided into two categories according to their different *C. difficile* infection/colonization statuses according to the criteria of previous studies^84^: (1) CD+ (n = 12): samples that displayed a positive result in clinical fecal nucleic acid amplification testing (Nucleic Acid Amplification Tests, NAAT; Xpert *C. difficile*/Epi) result. (2) CD− (n = 7): NAAT-negative stool on clinical *C. difficile* testing. These two categories of samples were utilized to design corresponding 16S rRNA gene primers using KuafuPrimer.

#### Validation cohort

Furthermore, a subsequent cohort of stool samples procured from patients exhibiting increased diarrhea or alterations in stool characteristics between September and November 2023 at the First Affiliated Hospital of Zhejiang University School of Medicine in Hangzhou, China, were employed to experimentally validate the designed primers (CD+, n=17; CD−, n = 17). Samples from patients under 18 years old or those who had undergone a colostomy were excluded. Additionally, samples from patients who had received antibiotics within the past 7 days were also excluded.

### DNA extraction

DNA was extracted from 0.5 g of stool material following the manufacturer’s instructions utilizing the OMG-Soil PF Mag-Bind Soil DNA Kit (Omega Bio-Tek, Georgi, USA). The quality and concentration of the extracted DNA were assessed by 1.0 % agarose gel electrophoresis and a NanoDrop 2000 spectrophotometer (Thermo Scientific Inc., United States), respectively. Samples with sufficient DNA concentration for library preparation were eluted in 70 μL elution solution C6 and kept at −80°C until use.

### Shotgun metagenomic sequencing

Samples that met the criteria for library construction were fragmented to an average length of approximately 400 bp utilizing Covaris M220 (Gene Company Limited, China), and subsequently shotgun metagenomic paired-end libraries were constructed using NEXTFLEX Rapid DNA-Seq (Bioo Scientific, Austin, TX, USA) in accordance with the manufacturer’s instructions. 2 × 150 bp paired-end sequencing was performed on Illumina NovaSeq™ X Plus instrument (Illumina Inc., San Diego, CA, USA) with at least 1,000,000 raw reads per sample.

### Analysis of shotgun metagenomic data

Fastp (version 0.23.4) was used to conduct quality control for metagenomic raw reads with the --qualified_quality_phred parameter set as 20, --length_required as 85 to remove reads of low quality or short length. And --low_complexity_filter parameter was set to eliminate reads of low complexity (i.e. long repetitive fragments). After that, Bowtie2 (version 2.3.4.1) was used to map reads to the human reference genome (GRCh38) to remove host sequences. Finally, Kraken2 (version 2.1.3) and Braken (version 2.9) were employed to taxonomically classify clean reads after host removal with standard Kraken 2 database (https://benlangmead.github.io/aws-indexes/k2).

### 16S amplicon sequencing

Only stool samples of the validation cohort were utilized for 16S amplicon sequencing. The universal primer pair 338F-806R (5′-ACTCCTACGGGAGGCAGCA-3′ and 5′-GGAC-TACHVGGGTWTCTAAT-3′) was used to amplify the V3-V4 region of the bacterial 16S rRNA gene for all samples. Additionally, KuafuPrimer-designed primers for CD+ (5′-GCCAGCAGCYGCGGTRAHAC-3′ and 5′-TTGTGCGGGBCCCCGTCWAT-3′) and CD− group (5′-GCTAAHTNYGTGCCAGCAGC-3′ and 5′-CCCCGTCWATTYHTTTGAGT-3′) were used to amplify the V4-V5 region of the bacterial 16S rRNA gene, respectively. All PCR were done using ABI GeneAmp® 9700 instrument (BIO-RAD, USA), Pro Taq 20μL reaction buffer with 27 cycles, an annealing temperature of 55°C and an elongation time of 30 sec. 16S rRNA gene libraries were constructed using the NEXTFLEX Rapid DNA-Seq Kit, and sequenced on an Illumina Nextseq 2000 (2 × 300 bp, Illumina, Hayward, California, USA) as described by the manufacturer.

### Analysis of 16S rRNA gene sequencing data

Fastp (version 0.23.4) was used to conduct quality control for 16S amplicon sequencing raw reads with the --qualified_quality_phred parameter set as 20, --cut_tail, and --low_complexity_filter parameters set as True. FLASH (version 1.2.11) was employed for the merge of paired-end reads. Vsearch (version 2.21.0) was used to remove chimeras. After that, the merged reads were mapped to the human reference genome (GRCh38) using Bowtie2 (version 2.3.4.1) with the --very-sensitive option to assess the ratio of off-target amplification of human DNA. Finally, Kraken2 (version 2.1.3) and Braken (version 2.9) were utilized for taxonomic assignment of clean reads with the SILVA SSU Ref NR 99 (release 138.1) database^79^. Each sample was rarefied to a uniform depth of 20,000 reads.

### Statistical analysis and visualization

Chao1 indices, Bray-Curtis similarities and Pearson correlation coefficients were calculated in Python using the skbio, scipy, and numpy packages. The accuracy and recall value of the universal primers and KuafuPrimer-designed primers were calculated in Python using scikit-learn package with metagenomic genus profiles as golden label. Statistical comparison was conducted using two-sided Wilcoxon signed-rank and rank-sum tests for paired and non-paired comparisons respectively, and a *p*-value_<_0.05 was considered significant for all analyses. Change points of KuafuPrimer improvement over universal primer in different environments were calculated in R using changepoint package. The entropy of conserved region was calculated in Python using scipy package. The visualization of data was performed in R (version 4.4.1) and Python (version 3.8).

## Data availability

The raw data for the gut samples have been deposited in the NCBI database under accession number PRJNA1150379. Additional data are available in the supplementary files and at https://github.com/zhanghaoyu9931/KuafuPrimer.

## Code availability

The code for KuafuPrimer in this paper is available at https://github.com/zhanghaoyu9931/KuafuPrimer.

## Author information

### Contributions

HQZ and XQJ conceived the study. HQZ, YHX, and XQJ supervised the work. HYZ and XQJ carried out the algorithm design and performed the result analysis. HYZ, XQJ, XWY, HYW, JHH, QG, SFW, HCY, XPG and JYG curated the data and plotted the figures, XQJ, HYZ, HQZ, AJ and ZW wrote and revised the manuscript. YHX, TTX and PL collected the gut samples. All authors read and accepted the final version of the manuscript.

## Acknowledgments

Part of the analysis was performed on the High-Performance Computing Platform of Peking University, and Biomedical Computing Platform of National Biomedical Imaging Center of Peking University. The authors thank Weiming Zhang and Shan Qiu for their valuable comments on the revision of this manuscript. We acknowledge Biorender (www.biorender.com) for providing the mapping platform.

## Ethics declarations

### Ethics approval and consent to participate

This study was approved by the ethics committee of the First Affiliated Hospital of Zhejiang University School of Medicine in Hangzhou, China (No: 2016-458-1).

### Consent for publication

Not applicable.

## References

1. Banerjee, S. & van der Heijden, M.G.A. Soil microbiomes and one health. Nat. Rev. Microbiol. 21, 6–20 (2023).

2. Gilbert, J.A. & Hartmann, E.M. The indoors microbiome and human health. Nat. Rev. Microbiol. 22, 742–755 (2024).

3. Riquelme, E. et al. Tumor microbiome diversity and composition influence pancreatic cancer outcomes. Cell 178, 795–806.e712 (2019).

4. The Integrative Human Microbiome Project. Nature 569, 641–648 (2019).

5. Lebeer, S. et al. A citizen-science-enabled catalogue of the vaginal microbiome and associated factors. Nat. Microbiol. 8, 2183–2195 (2023).

6. Shen, X. et al. Nonlinear dynamics of multi-omics profiles during human aging. Nature Aging 4, 1619–1634 (2024).

7. Zhou, X. et al. Longitudinal profiling of the microbiome at four body sites reveals core stability and individualized dynamics during health and disease. Cell Host Microbe 32, 506–526.e509 (2024).

8. Zhou, W. et al. Longitudinal multi-omics of host-microbe dynamics in prediabetes. Nature 569, 663–671 (2019).

9. Tebani, A. et al. Integration of molecular profiles in a longitudinal wellness profiling cohort. Nat. Commun. 11, 4487 (2020).

10. Lloyd-Price, J. et al. Multi-omics of the gut microbial ecosystem in inflammatory bowel diseases. Nature 569, 655–662 (2019).

11. Tito, R.Y. et al. Microbiome confounders and quantitative profiling challenge predicted microbial targets in colorectal cancer development. Nat. Med. 30, 1339–1348 (2024).

12. Xiao, L., Zhang, F. & Zhao, F. Large-scale microbiome data integration enables robust biomarker identification. Nat. Comput. Sci. 2, 307–316 (2022).

13. Park, C., Kim, S.B., Choi, S.H. & Kim, S. Comparison of 16S rRNA gene based microbial profiling using five next-generation sequencers and various primers. Front. Microbiol. 12, 715500 (2021).

14. Karst, S.M. et al. Retrieval of a million high-quality, full-length microbial 16S and 18S rRNA gene sequences without primer bias. Nat. Biotechnol. 36, 190–195 (2018).

15. Eloe-Fadrosh, E.A., Ivanova, N.N., Woyke, T. & Kyrpides, N.C. Metagenomics uncovers gaps in amplicon-based detection of microbial diversity. Nat. Microbiol. 1, 15032 (2016).

16. Zhang, R.Y. et al. Design of targeted primers based on 16S rRNA sequences in meta-transcriptomic datasets and identification of a novel taxonomic group in the Asgard archaea. BMC Microbiol. 20, 25 (2020).

17. Darwish, N., Shao, J., Schreier, L.L. & Proszkowiec-Weglarz, M. Choice of 16S ribosomal RNA primers affects the microbiome analysis in chicken ceca. Sci. Rep. 11, 11848 (2021).

18. Anantharaman, K. et al. Thousands of microbial genomes shed light on interconnected biogeochemical processes in an aquifer system. Nat. Commun. 7, 13219 (2016).

19. Chen, Z. et al. Impact of preservation method and 16S rRNA hypervariable region on gut microbiota profiling. mSystems 4, e00271–00218 (2019).

20. Graspeuntner, S., Loeper, N., Künzel, S., Baines, J.F. & Rupp, J. Selection of validated hypervariable regions is crucial in 16S-based microbiota studies of the female genital tract. Sci. Rep. 8, 9678 (2018).

21. Deissová, T. et al. 16S rRNA gene primer choice impacts off-target amplification in human gastrointestinal tract biopsies and microbiome profiling. Sci. Rep. 13, 12577 (2023).

22. Abellan-Schneyder, I. et al. Primer, pipelines, parameters: issues in 16S rRNA gene sequencing. mSphere 6, e01202–01220 (2021).

23. Breitenwieser, F., Doll, E.V., Clavel, T., Scherer, S. & Wenning, M. Complementary use of cultivation and high-throughput amplicon sequencing reveals high biodiversity within raw milk microbiota. Front. Microbiol. 11, 1557 (2020).

24. Takahashi, S., Tomita, J., Nishioka, K., Hisada, T. & Nishijima, M. Development of a prokaryotic universal primer for simultaneous analysis of bacteria and archaea using next-generation sequencing. PLoS ONE 9, e105592 (2014).

25. Walker, S.P. et al. Non-specific amplification of human DNA is a major challenge for 16S rRNA gene sequence analysis. Sci. Rep. 10, 16356 (2020).

26. Beckers, B. et al. Performance of 16S rDNA primer pairs in the study of rhizosphere and endosphere bacterial microbiomes in metabarcoding studies. Front. Microbiol. 7, 650 (2016).

27. Bedarf, J.R. et al. Much ado about nothing? Off-target amplification can lead to false-positive bacterial brain microbiome detection in healthy and Parkinson’s disease individuals. Microbiome 9, 75 (2021).

28. Wang, Y. & Qian, P.Y. Conservative fragments in bacterial 16S rRNA genes and primer design for 16S ribosomal DNA amplicons in metagenomic studies. PLoS ONE 4, e7401 (2009).

29. Johnson, J.S. et al. Evaluation of 16S rRNA gene sequencing for species and strain-level microbiome analysis. Nat. Commun. 10, 5029 (2019).

30. Kameoka, S. et al. Benchmark of 16S rRNA gene amplicon sequencing using Japanese gut microbiome data from the V1-V2 and V3-V4 primer sets. BMC Genomics 22, 527 (2021).

31. Regueira-Iglesias, A. et al. In silico evaluation and selection of the best 16S rRNA gene primers for use in next-generation sequencing to detect oral bacteria and archaea. Microbiome 11, 58 (2023).

32. Caporaso, J.G. et al. Global patterns of 16S rRNA diversity at a depth of millions of sequences per sample. Proc. Natl. Acad. Sci. U. S. A. 108 Suppl 1, 4516–4522 (2011).

33. Kim, S.W. et al. Robustness of gut microbiota of healthy adults in response to probiotic intervention revealed by high-throughput pyrosequencing. DNA Res. 20, 241–253 (2013).

34. Xie, N.G. et al. Designing highly multiplex PCR primer sets with Simulated Annealing Design using Dimer Likelihood Estimation (SADDLE). Nat. Commun. 13, 1881 (2022).

35. Untergasser, A. et al. Primer3--new capabilities and interfaces. Nucleic Acids Res. 40, e115 (2012).

36. Hugerth, L.W. et al. DegePrime, a program for degenerate primer design for broad-taxonomic-range PCR in microbial ecology studies. Appl. Environ. Microbiol. 80, 5116–5123 (2014).

37. Sambo, F. et al. Optimizing PCR primers targeting the bacterial 16S ribosomal RNA gene. BMC Bioinformatics 19, 343 (2018).

38. Jeon, H. et al. MRPrimerW2: an enhanced tool for rapid design of valid high-quality primers with multiple search modes for qPCR experiments. Nucleic Acids Res. 47, W614–w622 (2019).

39. Bae, J., Jeon, H. & Kim, M.S. GPrimer: a fast GPU-based pipeline for primer design for qPCR experiments. BMC Bioinformatics 22, 220 (2021).

40. Xia, H., et al. MultiPrime: A reliable and efficient tool for targeted nextlJgeneration sequencing. iMeta 2, e143 (2023).

41. Wang, L. & Jiang, T. On the complexity of multiple sequence alignment. J. Comput. Biol. 1, 337–348 (1994).

42. Yang, B., Wang, Y. & Qian, P.Y. Sensitivity and correlation of hypervariable regions in 16S rRNA genes in phylogenetic analysis. BMC Bioinformatics 17, 135 (2016).

43. Pascoal, F., Costa, R. & Magalhães, C. The microbial rare biosphere: current concepts, methods and ecological principles. FEMS Microbiol. Ecol. 97 (2021).

44. SantaLucia, J., Jr. & Hicks, D. The thermodynamics of DNA structural motifs. Annu. Rev. Biophys. Biomol. Struct. 33, 415–440 (2004).

45. Hartmann, M., Howes, C.G., Abarenkov, K., Mohn, W.W. & Nilsson, R.H. V-Xtractor: an open-source, high-throughput software tool to identify and extract hypervariable regions of small subunit (16S/18S) ribosomal RNA gene sequences. J. Microbiol. Methods 83, 250–253 (2010).

46. Stroup, E.K. & Ji, Z. Deep learning of human polyadenylation sites at nucleotide resolution reveals molecular determinants of site usage and relevance in disease. Nat. Commun. 14, 7378 (2023).

47. Kim, H. et al. MRPrimer: a MapReduce-based method for the thorough design of valid and ranked primers for PCR. Nucleic Acids Res. 43, e130 (2015).

48. Lott, M.T. et al. mtDNA variation and analysis using Mitomap and Mitomaster. Curr Protoc Bioinformatics 44, 1.23.21–26 (2013).

49. Song, L. & Xie, K. Engineering CRISPR/Cas9 to mitigate abundant host contamination for 16S rRNA gene-based amplicon sequencing. Microbiome 8, 80 (2020).

50. Wang, F. et al. Assessment of 16S rRNA gene primers for studying bacterial community structure and function of aging flue-cured tobaccos. AMB Express 8, 182 (2018).

51. Klindworth, A. et al. Evaluation of general 16S ribosomal RNA gene PCR primers for classical and next-generation sequencing-based diversity studies. Nucleic Acids Res. 41, e1 (2013).

52. Zhang, J. et al. Evaluation of different 16S rRNA gene V regions for exploring bacterial diversity in a eutrophic freshwater lake. Sci. Total Environ. 618, 1254–1267 (2018).

53. Shi, S., Kumar, S., Young, S., Maclean, P. & Jauregui, R. Evaluation of 16S rRNA gene primer pairs for bacterial community profiling in an across soil and ryegrass plant study. Journal of Sustainable Agriculture and Environment 2, 500–512 (2023).

54. Chen, Y., Knight, R. & Gallo, R.L. Evolving approaches to profiling the microbiome in skin disease. Front. Immunol. 14, 1151527 (2023).

55. Meisel, J.S. et al. Skin microbiome surveys are strongly influenced by experimental design. J. Invest. Dermatol. 136, 947–956 (2016).

56. Thomas, F. et al. Evaluation of a new primer combination to minimize plastid contamination in 16S rDNA metabarcoding analyses of alga-associated bacterial communities. Environ. Microbiol. Rep. 12, 30–37 (2020).

57. Liu, C. et al. Soil bacterial communities of three types of plants from ecological restoration areas and plant-growth promotional benefits of Microbacterium invictum (strain X-18). Front. Microbiol. 13, 926037 (2022).

58. Shahzad, A. et al. Heavy metals mitigation and growth promoting effect of endophytic Agrococcus terreus (MW 979614) in maize plants under zinc and nickel contaminated soil. Front. Microbiol. 14, 1255921 (2023).

59. Ribeiro, I.D.A. et al. Antifungal potential against Sclerotinia sclerotiorum (Lib.) de Bary and plant growth promoting abilities of Bacillus isolates from canola (Brassica napus L.) roots. Microbiol. Res. 248, 126754 (2021).

60. Cordovez, V. et al. Priming of plant growth promotion by volatiles of root-associated Microbacterium spp. Appl. Environ. Microbiol. 84 (2018).

61. Teramoto, K. et al. Classification of Cutibacterium acnes at phylotype level by MALDI-MS proteotyping. Proc. Jpn. Acad. Ser. B Phys. Biol. Sci. 95, 612–623 (2019).

62. de Assis, D.A., Machado, C., Matte, C. & Ayub, M.A.Z. High cell density culture of dairy Propionibacterium sp. and Acidipropionibacterium sp.: a review for food industry applications. Food Bioproc. Tech. 15, 734–749 (2022).

63. Traisaeng, S. et al. Leuconostoc mesenteroides fermentation produces butyric acid and mediates Ffar2 to regulate blood glucose and insulin in type 1 diabetic mice. Sci. Rep. 10, 7928 (2020).

64. Geerlings, S.Y., Kostopoulos, I., de Vos, W.M. & Belzer, C. Akkermansia muciniphila in the human gastrointestinal tract: when, where, and how? Microorganisms 6, 75 (2018).

65. Belzer, C. & de Vos, W.M. Microbes inside--from diversity to function: the case of Akkermansia. ISME J. 6, 1449–1458 (2012).

66. Bae, M. et al. Akkermansia muciniphila phospholipid induces homeostatic immune responses. Nature 608, 168–173 (2022).

67. Zhao, Y., Zhu, Z., Chen, X. & Li, Y. Discovery of a novel potential polyphosphate accumulating organism without denitrifying phosphorus uptake function in an enhanced biological phosphorus removal process. Sci. Total Environ. 912, 168952 (2024).

68. Zhao, S., Rogers, M.J. & He, J. Abundance of organohalide respiring bacteria and their role in dehalogenating antimicrobials in wastewater treatment plants. Water Res. 181, 115893 (2020).

69. Ji, B., Liu, C., Liang, J. & Wang, J. Seasonal succession of bacterial communities in three eutrophic freshwater lakes. Int. J. Environ. Res. Public. Health 18, 6950 (2021).

70. Zhang, Y. et al. Long-term effects of three compound probiotics on water quality, growth performances, microbiota distributions and resistance to Aeromonas veronii in crucian carp Carassius auratus gibelio. Fish Shellfish Immunol. 120, 233–241 (2022).

71. Shi, T., Wang, Y.F., Wang, H. & Wang, B. Genus Nocardiopsis: a prolific producer of natural products. Mar. Drugs. 20 (2022).

72. Mihai, P. et al. A framework for human microbiome research. Nature 486, 215–221 (2012).

73. Wang, G. et al. Ginsenoside Rg3 enriches SCFA-producing commensal bacteria to confer protection against enteric viral infection via the cGAS-STING-type I IFN axis. ISME J. 17, 2426–2440 (2023).

74. Ke, S. et al. Integrating gut microbiome and host immune markers to understand the pathogenesis of Clostridioides difficile infection. Gut Microbes 13, 1–18 (2021).

75. Ranjan, R., Rani, A., Metwally, A., McGee, H.S. & Perkins, D.L. Analysis of the microbiome: advantages of whole genome shotgun versus 16S amplicon sequencing. Biochem. Biophys. Res. Commun. 469, 967–977 (2016).

76. Brumfield, K.D., Huq, A., Colwell, R.R., Olds, J.L. & Leddy, M.B. Microbial resolution of whole genome shotgun and 16S amplicon metagenomic sequencing using publicly available NEON data. PLoS ONE 15, e0228899 (2020).

77. Thijs, S. et al. Comparative evaluation of four bacteria-specific primer pairs for 16S rRNA gene surveys. Front. Microbiol. 8, 494 (2017).

78. Thomas, M.C., Thomas, D.K., Selinger, L.B. & Inglis, G.D. spyder, a new method for in silico design and assessment of 16S rRNA gene primers for molecular microbial ecology. FEMS Microbiol. Lett. 320, 152–159 (2011).

79. Quast, C. et al. The SILVA ribosomal RNA gene database project: improved data processing and web-based tools. Nucleic Acids Res. 41, D590–596 (2013).

80. Martinez-Porchas, M., Villalpando-Canchola, E., Ortiz Suarez, L.E. & Vargas-Albores, F. How conserved are the conserved 16S-rRNA regions? PeerJ 5, e3036 (2017).

81. Ronneberger, O., Fischer, P. & Brox, T. in Medical image computing and computer-assisted intervention–MICCAI 2015: 18th international conference, Munich, Germany, October 5-9, 2015, proceedings, part III 18 234–241 (Springer, 2015).

82. Edgar, R.C. MUSCLE: multiple sequence alignment with high accuracy and high throughput. Nucleic Acids Res. 32, 1792–1797 (2004).

83. Wang, K. et al. MFEprimer-3.0: quality control for PCR primers. Nucleic Acids Res. 47, W610–w613 (2019).

84. Berkell, M. et al. Microbiota-based markers predictive of development of Clostridioides difficile infection. Nat. Commun. 12, 2241 (2021).

