## Supplementary for "KuafuPrimer: Machine learning empowers the design of 16S amplicon sequencing primers toward minimal bias for bacterial communities"

**Extended Data Figures**

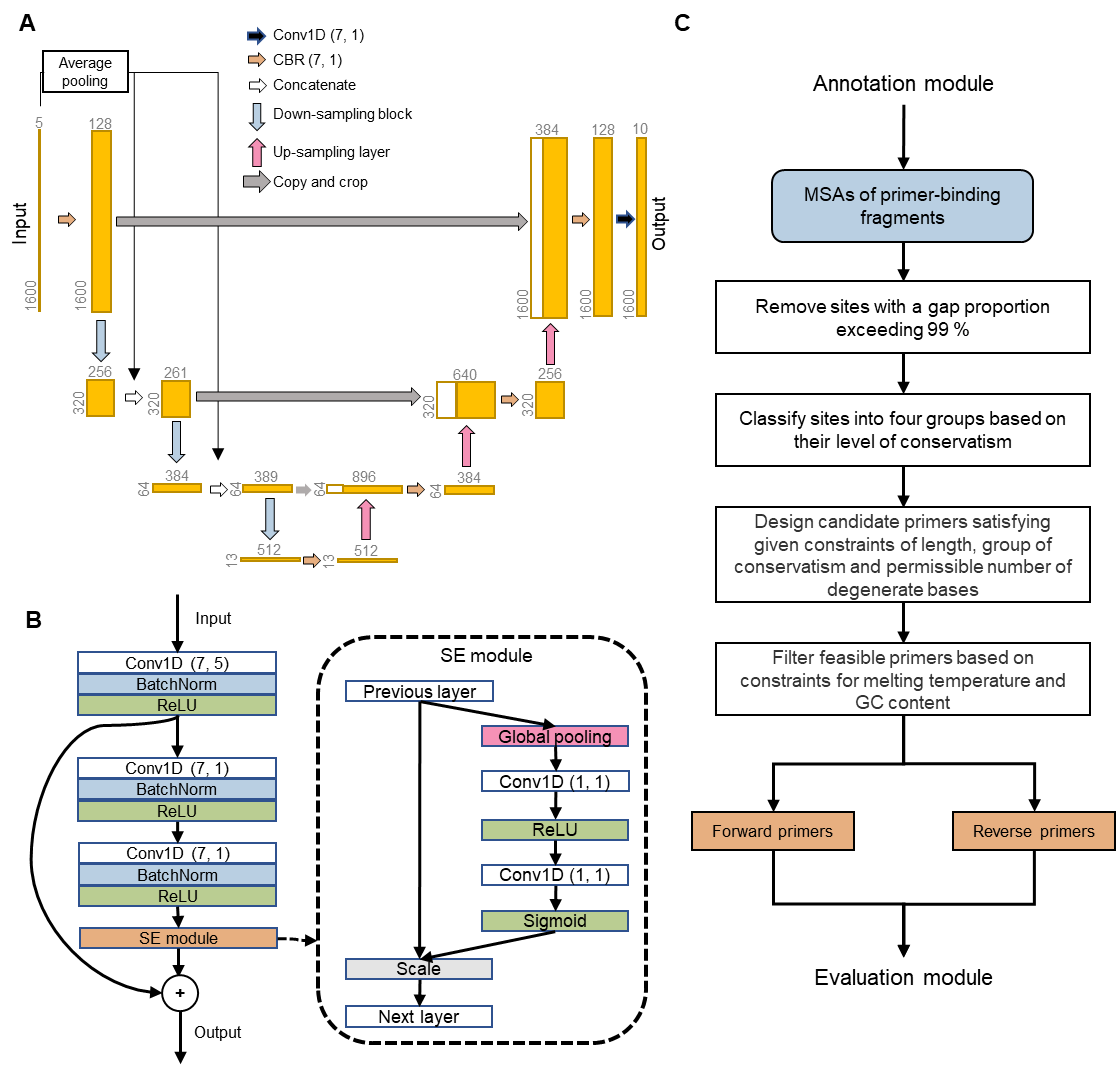

**Extended Data Fig. 1.** **The detailed structure of KuafuPrimer. A**. The U-shaped structure of DeepAnno16. Conv1D: one-dimensional convolutional neural network layer; CBR: a module comprising a Conv1D layer, a batch normalization (BatchNorm) layer, and a ReLU activation layer. The numbers in Conv1D layer and CBR module indicate the kernel size and stride, respectively. The up-sampling layer uses the nearest-neighbor interpolation method with a scale factor of five. **B**. The structure of the down-sampling block. The scale layer in the Squeeze-and-Excitation (SE) module applies channel-wise importance factors (output from a sigmoid layer) to the corresponding channel values. **C**. Workflow of the fast primer design module in KuafuPrimer.

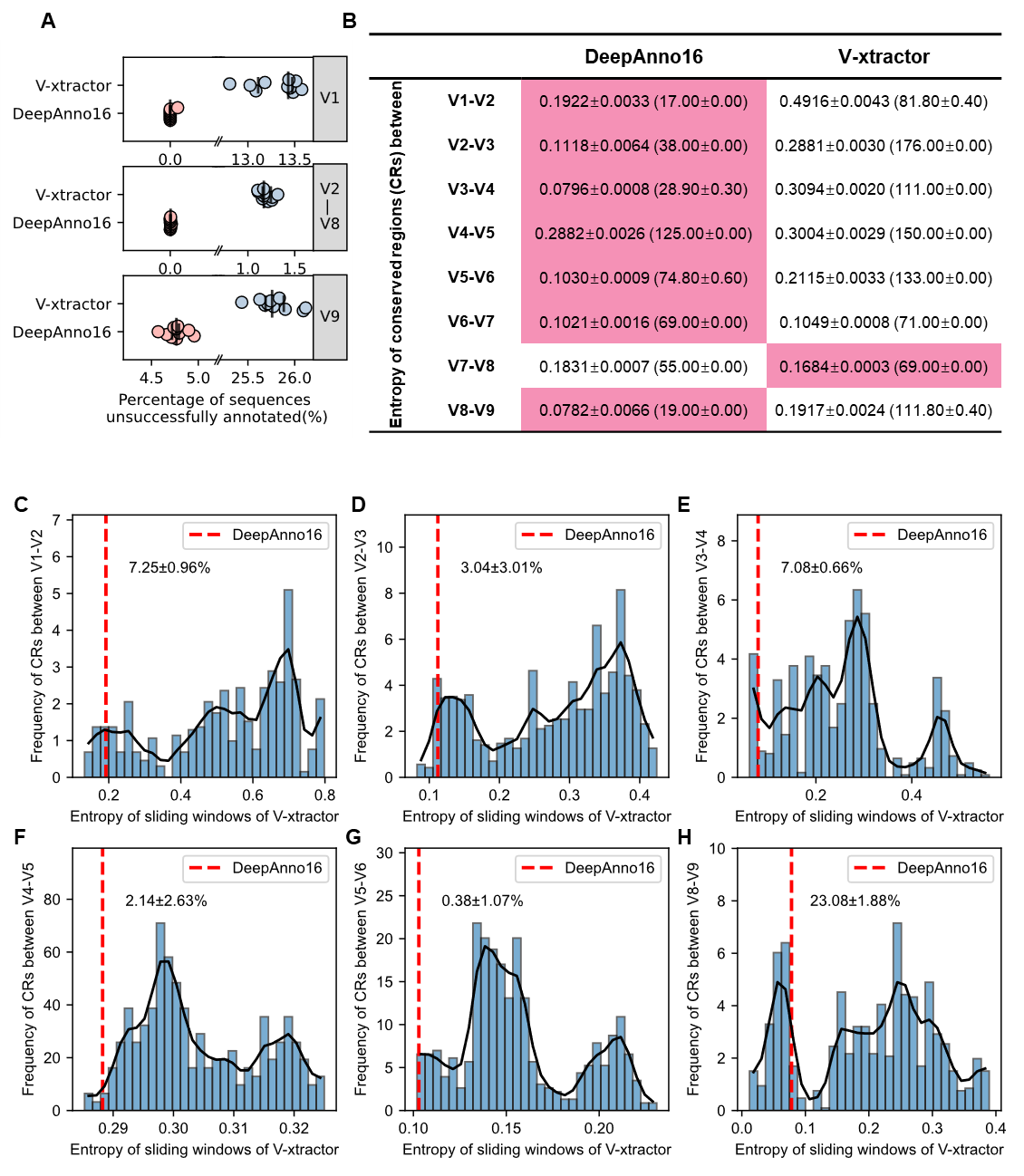

**Extended Data Fig. 2. Performance comparison of DeepAnno16 and V-xtractor. A**. Comparison of the percentage of sequences that failed to be annotated by DeepAnno16 and V-xtractor across different V-regions. **B**. Comparison of the entropy of the conserved regions (CRs) annotated by DeepAnno16 and V-xtractor. Entropy, a commonly used metric for site variability, is inversely correlated with conservatism (lower entropy indicates higher conservatism). The mean entropy values (± standard deviation) from 10-fold cross validations are shown, with the lengths of the CRs indicated in parentheses. Values highlighted in red denote a significant advantage relative to the competing tool (*p* < 0.05, Wilcoxon signed-rank test). **C-H**. Comparison of entropy distributions for CRs of identical lengths annotated by DeepAnno16 and V-xtractor. CRs annotated by DeepAnno16 are generally shorter than those annotated by V-xtractor (Extended Data Fig. S2B). CRs between V6-V7 and V7-V8 are not included due to their comparable lengths as annotated by the two tools. To mitigate the potential impact of CR­ length on entropy, sequences matching the lengths of DeepAnno16 annotations were extracted from V-xtractor annotations using a sliding window approach. The fitted curve of the distribution is depicted with a black line. Red dashed vertical lines indicate the entropy values of the CRs annotated by DeepAnno16, with the adjacent numbers representing the proportion of sliding window CRs annotated by V-xtractor that exhibit lower entropy values. Sequences that could not be successfully annotated were excluded, which may have led to an overestimation of conservatism for V-xtractor annotations particularly in the CRs between V8-V9, where V-xtractor exhibited low success rates.

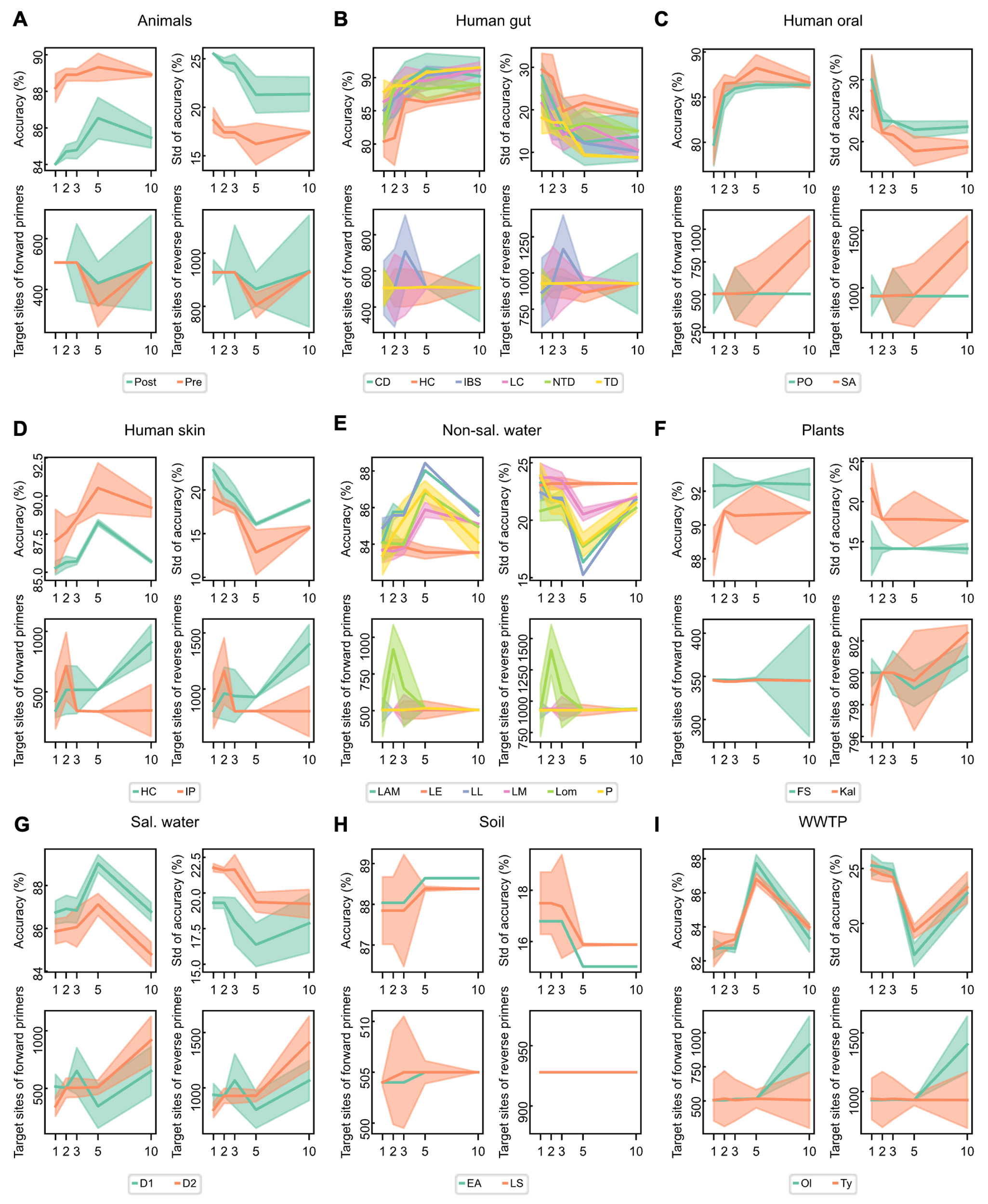

**Extended Data Fig. 3.** **Performance of primers designed by KuafuPrimer as the number of metagenomic samples used for training increases.** The mean taxonomic accuracy (top left), standard deviation of taxonomic accuracy for all genera (top right), and target sites of forward (bottom left) and reverse (bottom right) primers designed by KuafuPrimer based on different numbers of training samples (the x-axis) in each group (**A-I**), with ten repetitions. Solid line: median of ten repetitions. Transparent area: range between the upper and lower quartiles of the ten repetitions. Std: standard deviation.

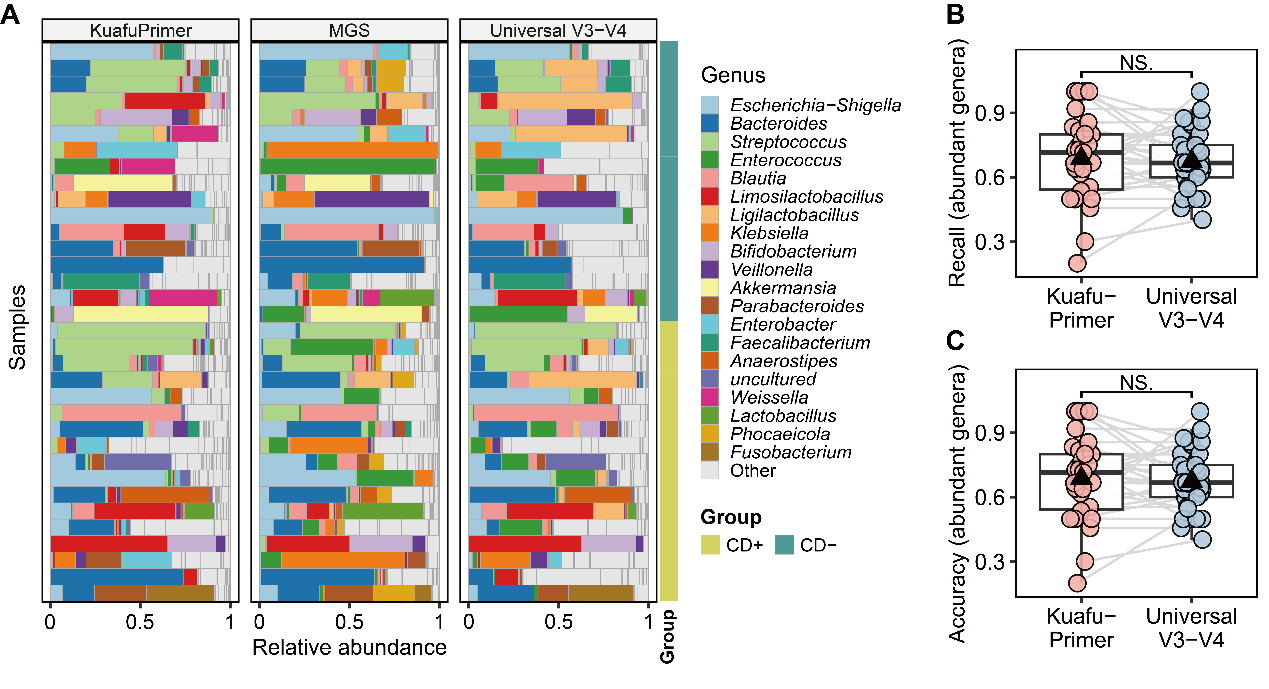

**Extended Data Fig. 4. Experimental results of KuafuPrimer****-designed primers, the** **universal V3-V4 primer and metagenomic sequencing.** **A**. Relative abundances of top 20 genera in 34 human gut samples as determined by KuafuPrimer-designed primers, the universal V3–V4 primer, and metagenomic sequencing. Using the genera identified in metagenomic samples as the gold standard, we further calculated the recall (**B**) and accuracy (**C**) of abundant genera for both KuafuPrimer-designed primers and the universal V3–V4 primer. NS.: Not significant, Wilcoxon signed-rank test. CD+: CDI positive group, n = 17; CD-: CDI negative group, n = 17.

**Supplementary methods**

**Annotation of** **16S rRNA gene sequences**

Primer contigs were aligned to each 16S rRNA gene sequence using BLAST (version 2.6.0+), with parameters -word_size 4 and -evalue 100. Aligned regions were annotated as conserved regions, while unaligned regions were identified as corresponding V-regions. Primer contigs were collected from previous studies (Supplementary Table S9). Sequences with any mismatched primer contigs (i.e., no hit or a hit of insufficient length) or incorrect alignment were excluded.

**KuafuPrimer algorithm**

Annotation module: The input nucleotide sequence for DeepAnno16 is first encoded into a “one-hot” vector $x_{1}\ldots x_{L}$, where $x_{i}\mathbb{\in R}^{d}$ is a binary vector of length d = 5 (four for nucleotides and one for padding). In this encoding, only one bit is active at each position, representing the corresponding nucleotide type. Each sequence is either padded or clipped to a fixed length of 1600 bp, in line with the requirement of deep neural networks for uniform input sizes. A position-wise vector $\overset{⃑}{y}=y_{1}\ldots y_{L}$ of 10 classes is used as the gold standard label based on previous annotation results. Here, $y_{i}\mathbb{\in R}$indicates the V-region (V1-V9) or the conserved region that the *i*^th^ bp of the input sequence belongs to. DeepAnno16 employs a U-shaped encoder-decoder architecture, consisting of interconnected down-sampling and up-sampling blocks (Extended Data Fig. S1A). The encoder consists of three down-sampling blocks, which generate condensed, globally informative feature maps from the original input sequence. Each down-sampling block incorporates an SE-ResNet module, comprising three Conv1D-BatchNorm-ReLU (CBR) layers followed by a squeeze-and-excitation (SE) module and a residual connection (Extended Data Fig. S1B). To avoid the loss of spatial information during the down-sampling process, the low-resolution representations of the original sequence are attached to the output of each down-sampling block. The decoder is primarily composed of three up-sampling blocks, which restore the sequence length reduced by the down-sampling counterpart to predict annotations for each position in the input sequence. Each up-sampling block contains an up-sampling layer using the nearest-extrapolate method to restore the original sequence length, followed by a concatenate operator and a CBR block. The concatenate operator combines the feature map of the corresponding down-sampling block with the output of the up-sampling layer, and the CBR block further refines the restored information. DeepAnno16 is trained using a 10-fold cross-validation strategy. The dataset is split into 10 groups, and for each training process, one group is used as the testing set, while the remaining groups are divided into training and validation sets according to a ratio of 8:1. The training and validation sets are used are used to develop the model and tune hyperparameters (learning rate, batch size, dropout ratio and early stopping), while the testing set is used to evaluate the model's performance and generalization. The average performance across all testing sets is used for model comparison. Subsequently, DeepAnno16 outputs all possible V-regions and extracts the corresponding primer-binding fragments. For instance, when targeting the V3-V4 region, the primer-binding fragment is extracted from the rightmost V2 site to the leftmost V3 site for forward primer design, and from the rightmost V4 site to the leftmost V5 site for reverse primer design. To ensure the integrity of primer-binding regions, we extend the regions by a certain number (default: 50) of sites at both the left and right ends when extracting fragments.

Fast primer design module: Primers are designed based on multiple sequence alignments (MSAs) of primer-binding fragments through the following steps:

1. Any site in the MSAs with a gap proportion exceeding 99 % is removed. This typically indicates that most representative sequences did not align to this site, which could be due to high sequence heterogeneity or errors in the MSA.

2. Sites are categorized into one of four groups based on their level of conservatism: gap-type sites, where the proportion of gaps exceeds 5 %; conserved sites, where a single base type exceeds 99.5 % of occurrences; degenerate sites, which feature multiple base types each occurring at a proportion greater than 0.1 % and a combined frequency of them exceeding 99.5 %, with assigned degenerate bases based on the degenerate base table; and normal sites, all remaining sites that do not fall into the categories above.

3. Any consecutive sequence that meets length criteria (default: 18-22 bp), does not contain gap-type sites or more than three degenerate sites, and has at most one normal site (allowing for one site mismatch), will be included in the candidate primer set.

4. Finally, forward and reverse primers are filtered to ensure that they have a melting temperature between 55 ℃ and 65 ℃ and a GC content ranging from 0.4 to 0.6.

**Collection of universal primer pairs.** Through a comprehensive literature review, we identified seven universal primer pairs commonly used in 16S amplicon analysis (Supplementary Table 4). The amplicons generated by these primers range in length from approximately 300 to 500 bp, which is within the sequencing length constraints of next-generation sequencing (NGS) platforms. These primers have been widely employed in amplicon sequencing across diverse environments, making them suitable for the needs of most researchers.

In addition, nine Supplementary Table Files contain data that is too large to be displayed on a

single page:

**Supplementary Table S1** Metadata of 26 environmental and host-associated groups.

**Supplementary Table S2** Results of metagenomic analysis of each sample.

**Supplementary Table S3** Ratio of genera coveraged by different number of training samples for 26 environmental and host-associated groups.

**Supplementary Table S4** Universal primers used for *in silico* PCR comparison.

**Supplementary Table S5** Target sites and *in silico* PCR accuracy of primers designed by KuafuPrimer and universal primers across 26 environmental and host-associated groups.

**Supplementary Table S6** Taxonomic accuracy of primers designed by KuafuPrimer based on different number of training samples for HC (healthy control group, human gut).

**Supplementary Table S7** Metadata of longitudinal studies.

**Supplementary Table S8** Target sites and *in silico* PCR accuracy of primers designed by KuafuPrimer and universal primers for longitudinal studies.

**Supplementary Table S9** Primer contigs used for annotation of variable and conserved regions of 16S rRNA gene.

**Supplementary Table S10** Metadata of participants in CD+ and CD- groups.
